# Temporal Genomic Dynamics Shape Clinical Trajectory in Multiple Myeloma

**DOI:** 10.1101/2024.08.30.610457

**Authors:** Francesco Maura, Marcella Kaddoura, Alexandra M. Poos, Linda B. Baughn, Bachisio Ziccheddu, Marc-Andrea Bärtsch, Anthony Cirrincione, Kylee Maclachlan, Monika Chojnacka, Benjamin Diamond, Marios Papadimitriou, Patrick Blaney, Lukas John, Philipp Reichert, Stefanie Huhn, Dylan Gagler, Yanming Zhang, Ahmet Dogan, Alexander M Lesokhin, Faith Davies, Hartmut Goldschmidt, Roland Fenk, Katja C. Weisel, Elias K. Mai, Neha Korde, Gareth J Morgan, S. Vincent Rajkumar, Shaji Kumar, Saad Usmani, Ola Landgren, Marc S. Raab, Niels Weinhold

## Abstract

To comprehensively unravel the temporal relationship between initiating and driver events and its impact on clinical outcomes, we analyzed 421 whole-genome sequencing profiles from 382 patients. Using clock-like mutational signatures, we estimated a time lag of 2-4 decades between initiating events and diagnosis. In patients with hyperdiploidy, we demonstrate that trisomies of odd-numbered chromosomes can be acquired simultaneously with other chromosomal gains, such as 1q gain. We provide evidence that hyperdiploidy is acquired after canonical IGH translocation when both events are present. Finally, patients with early 1q gain had adverse outcomes similar to those with 1q amplification (>1 extra-copies), but faring worse than those with late 1q gain. This underscores that the prognostic impact of 1q gain/amp depends more on the timing of acquisition than on the number of extra copies gained. Overall, this study contributes to a better understanding of the life history of MM and may have prognostic implications.

## INTRODUCTION

The pathogenesis of multiple myeloma (MM) involves a protracted evolution through distinct asymptomatic precursor stages, including monoclonal gammopathy of undetermined significance (MGUS) and smoldering multiple myeloma (SMM)^1–6^. The initiation of this process is believed to occur within the germinal center (GC), where a B-cell acquires specific somatic events, such as hyperdiploidy (HY) or immunoglobulin heavy chain (IGH) translocations^7–17^. Historically, these events are recognized to have a key initiating role in MM pathogenesis due to their clonal nature from even the earliest disease phases and their major influence on gene expression^1,8,9,11,14,18–22^. HY, the most common event, is detectable in approximately 60% of MM cases and involves multiple trisomies of odd-numbered chromosomes (3, 5, 7, 9, 11, 15, 19, and 21)^3^. The second group comprises translocations between the IGH locus and various oncogenes, including *CCND1*, *MAF*, *MAFB*, and *NSD2* (i.e., canonical IGH translocations)^8,10,11^. Notably, approximately 10% of MM patients carry both a canonical IGH translocation and a HY profile^8,14,16,21,23^ and it remains unclear which of these two events occurs first (**Supplementary Data 1**).

Following these initiating events, the evolution to active MM involves the acquisition of additional genomic drivers such as aneuploidies, driver gene mutations and structural variants (SV)^21,24^. Among the latter, SV involving the *MYC* locus represent the second most frequent somatic event in MM, detectable in 40-50% of patients and enriched in HY^22,25,26^. Considering the low prevalence among MGUS and SMM, *MYC* SV are considered late events acquired after HY and during the progression from SMM to MM^19,20,22,27,28^. In addition to HY, canonical IGH translocations, and *MYC* SV, the gain of chromosome 1q (1q gain) has emerged as one of the most important MM somatic events. This copy number variant (CNV) is detectable in approximately 30-40% of newly diagnosed MM (NDMM) cases and is associated with a poor prognosis^16,29–31^. The presence of two or more extra copies of 1q (i.e., 1q amp) has been associated with worse prognosis compared to single 1q gain, but this has not been confirmed by most recent studies^32–34^. Despite its biological and clinical relevance, it remains largely unclear when 1q gain is acquired during the MM pathogenesis. While 1q gain can be detected in precursor conditions many years before disease progression, it may also manifest only at relapse^35,36^. Recent evidence suggests that the latter clinical scenario is consistent with the treatment-induced selection of a pre-existing subclone with 1q gain^37^. Nevertheless, the possibility that 1q gain may be acquired after treatment cannot be ruled out.

Several pieces of evidence suggest that the complex progression from initiation of MGUS to SMM and ultimately MM spans multiple decades^1,6,38–40^. Notably, diverse epidemiological findings show that conditions like MGUS and the presence of monoclonal proteins, detectable through mass spectrometry, can manifest 10-20 years before SMM or MM diagnosis^6,38,41^. Moreover, utilizing clock-like single base substitution (SBS) mutational signatures like SBS1 and SBS5^42^, it has been demonstrated on limited data set that the temporal gap between the initial large chromosomal duplications (e.g., HY) and MM diagnosis can extend beyond 30 years^35,43^. This substantial time lag could have a profound impact on tumor evolution and clinical behavior, given the diverse temporal patterns of genomic events. However, the limited samples size of these studies has hindered further investigations.

To decipher the temporal relationship between initiating and driver events and its impact on clinical outcomes, we interrogated a large series of high coverage (80X) MM whole genome sequencing (WGS, n=421). Leveraging molecular time workflow ^43–45^, our analysis indicates that HY is heterogeneously acquired during MM life history, earlier in the absence of IGH translocations and later when both events co-occur at time of diagnosis. For 1q gain, it may occur simultaneously with initiating trisomies in some HY MM cases, but can also be newly acquired after front line treatment. Furthermore, our findings demonstrate that the poor prognostic impact of 1q gain is influenced more so by its timing rather than the number of copies. Specifically, early 1q gain is associated with unfavorable clinical outcomes in NDMM. Overall, our data reveal that the temporal acquisition of distinct genomic drivers has a considerable effect on MM tumor evolution and clinical outcomes.

## RESULTS

### Multiple myeloma mutational signatures landscape

We interrogated 421 WGS from 382 patients with MM. 323 and 98 WGS were generated from samples collected at diagnosis and relapse, respectively. Among the 315 NDMM patients with available clinical and treatment data, 68 were treated with VRd (bortezomib, lenalidomide dexamethasone), 187 with elotuzumab (Elo) + VRd, 16 with KRd (carfilzomib, lenalidomide dexamethasone), and 44 with daratumumab (dara) + KRd, respectively^46–48^. Overall, 230 (73%) NDMM patients underwent high-dose melphalan, and autologous stem cell transplantation (HDM-ASCT). HY and canonical IGH translocations were detected in 233 (55%) and 185 (44%) samples, respectively. Key clinical data are summarized in **Supplementary Tables 1** and **2**.

Median mutational burden was 5,492 (range 685-71,372) with relapsed refractory (RR) MM having a significantly higher mutational burden than NDMM (p<0.0001 using Wilcoxon test). Interrogating the genomic driver landscape and the prevalence of the 12 recently reported genomic subgroups^33^ we did not find any differences between the German and MSKCC cohort of NDMM patients (**Supplementary Tables 3-4** and **Supplementary Fig. 1**). Moreover, no differences were observed in mutational burden between the two cohorts (median 4912 and 5178, in the German and MSKCC cohort, respectively, **Supplementary Fig. 2A**).

To investigate the mutational processes involved in MM life-history, we performed SBS signatures analysis^49–51^. Nine main SBS signatures caused by 7 mutational processes were identified: APOBEC (SBS2 and SBS13), aging (SBS1 and SBS5), poly-eta in the GC (SBS9), radical oxygen stress damage (ROS), SBS8 (unknown etiology), and exposure to melphalan (SBS-MM1/SBS99) and platinum (SBS31) (**Fig. 1A**; https://cancer.sanger.ac.uk/cosmic/signatures/SBS/)^52^. APOBEC mutational activity was found in 90.7% of samples, with 66 (15.7%) defined as hyper-APOBEC (i.e., combined SBS2 and SBS13 contribution ≥ 11%) ^33^. In line with prior findings, patients with hyper-APOBEC had poor outcomes despite being exposed to highly effective combinations (**Supplementary Fig. 2B-C**)^53,54^ Also consistent with previous studies, *MAF*/*MAFB* translocations were characterized by a hyper-APOBEC profile in 19/22 patients (86.3%; **Fig. 1A**) ^53–55^.

**Figure 1.**
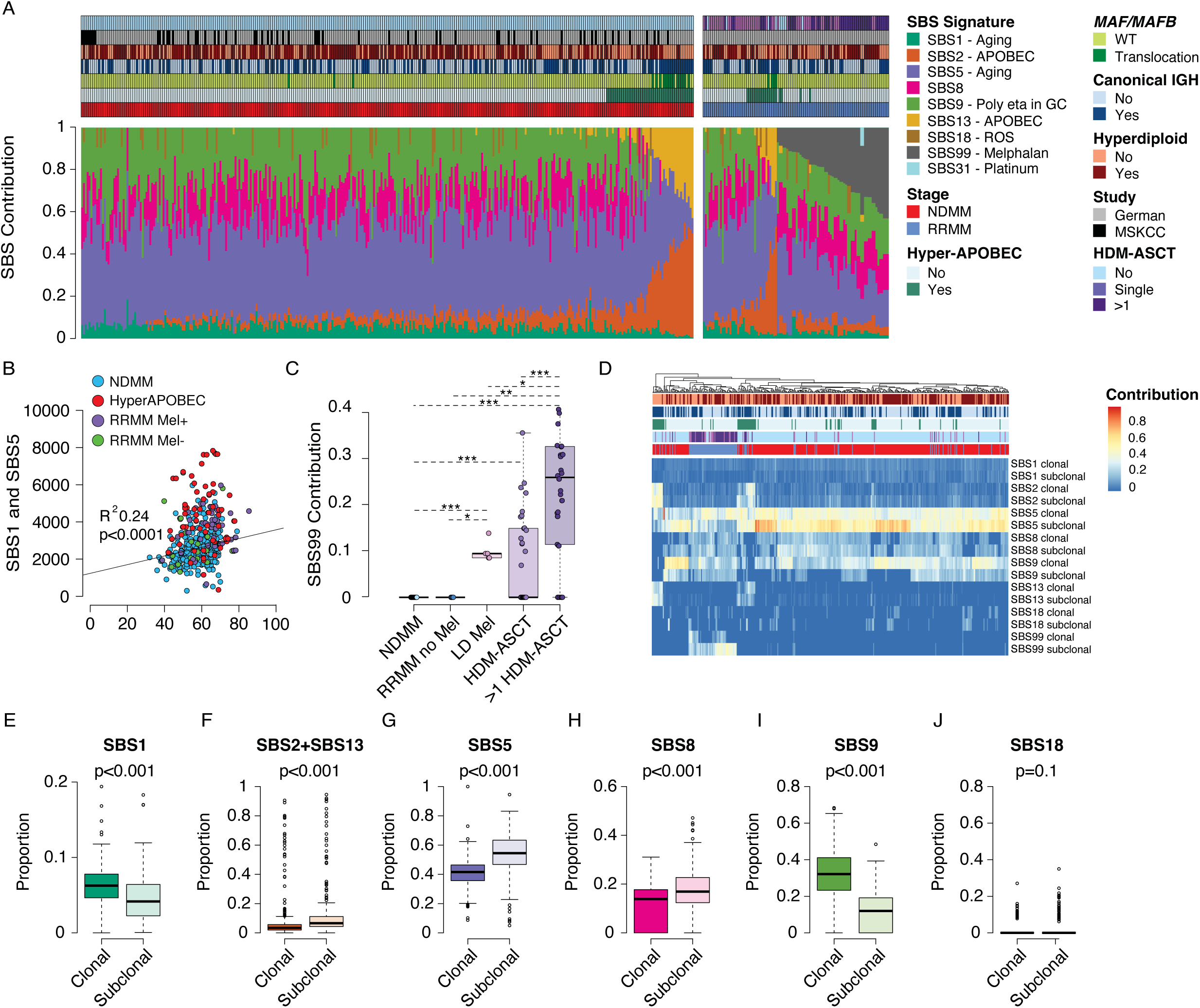
Multiple myeloma mutational signatures landscape. **A)** Contribution of SBS mutational signatures across 421 WGS from patients with MM. **B)** Correlation between SBS1 and SBS5 mutational burden (clock-like signatures) and patients’ age at sample collection. The p value and R^2^ were calculated using *lm* R function corrected for stage (relapsed/refractory vs newly diagnosed multiple myeloma) and presence of hyper-APOBEC. Three samples with more than 8000 SBS1 and SBS5 mutations were not included in the plot for graphical purpose (MM119, MM165, and RRMM48). **C)** Boxplots showing the SBS99 contribution per samples across stages and according to melphalan exposure. P-values were generated using Wilcoxon test ( ***: <0.001, **: 0.01; *: <0.05). Only p-values<0.05 are shown. **D)** SBS mutational signature contribution among clonal and subclonal variants per sample. Legend for the color scale is identical to Fig 1A. **E-J)** Comparison between clonal and subclonal variants for SBS mutational signature contribution. P-values were generated using a paired Wilcoxon test. SBS: single base substitution; GC: germinal center; ROS: radical oxidative stress; NDMM: newly diagnosed multiple myeloma; WT: wild type; RRMM: relapsed refractory multiple myeloma; HDM-ASCT: high dose melphalan and autologous stem cell transplantation; LD-Mel: low dose melphalan. Mel+ and Mel-: melphalan exposed or not exposed RRMM patients, respectively.

SBS1 and SBS5 were detected in all patients and were linearly correlated with age after correcting for stage and hyper-APOBEC (**Fig. 1A-B; Supplementary Data 2**). Forty-six samples (10.9%) did not fit into the clock-like SBS1-SBS5 linear model, with 25 (54.4%) having high APOBEC contribution and subsequent higher overall mutational burden. (i.e. hyper-APOBEC; **Supplementary Fig. 3A**). The diminished correlation observed between SBS1/SBS5 and age in individuals exhibiting hyper-APOBEC is consistent with findings in other tumors characterized by active hypermutators (e.g., UV-light, APOBEC, miss-match deficiency)^56^. This discrepancy is likely attributed to the inherent challenges faced by mutational signature tools in accurately quantifying mutational processes beyond the dominant hypermutator mechanism. Although their relative contributions differ, SBS1 and SBS5 are known to act like a molecular clock in most cancers and in virtually all normal human and mammalian tissues^42,62–64^. This molecular clock has been shown to be independent of replication rate, and conserved both in non-replicating normal tissues such as neurons ^62,65^, and highly replicating tumors such as Burkitt lymphoma ^44^. The SBS1 and SBS5 mutational rate was similar in our set of NDMM and in B-cell lymphomas (**Supplementary Fig. 3B**). Importantly, the mutation rate of SBS1 and SBS5 clonal variants observed in NDMM without hyper-APOBEC was similar to that reported for single-cell-colony expansions of normal memory B-cells (**Supplementary Fig. 2C; Supplementary Data 2**), supporting the existence of the SBS1-SBS5 molecular clock in most MM patients.

SBS9 was the third most frequent mutational signature, undetectable in only 38 (9%) patients, 37 (97%) of whom were hyper-APOBEC. SBS8 and SBS18 were detected in 340 (80.7%) and 47 (11.6%) patients, respectively. Only two patients had evidence of SBS31, both relapsed/refractory MM (RRMM) previously exposed to platinum-based regimens. SBS99 (previously reported as SBS-MM1 or Signature R) was observed in 59 (14%) samples, all collected from RRMM. All these patients were exposed to either HDM-ASCT (n=54) or low dose melphalan (n=5) ^36,57,58^. Overall, among 89 RRMM exposed to one (n=45) or more than one HDM-ASCT (n=44), 60% had evidence of SBS99. Interestingly, patients exposed to tandem HDM-ASCT (n=36) had higher SBS99 contribution compared to those who underwent single HDM-ASCT (n=18; p<0.001 using Wilcoxon test; **Fig. 1C**). This is the first evidence of cumulative mutagenic impacts of melphalan in MM. Five patients were positive for SBS99 after being exposed to low dose melphalan (i.e., bortezomib-melphalan-prednisone), confirming the mutagenic activity of this alkylating agent used in non-myeloablative doses^59^.

To investigate the temporal evolution of each SBS mutational signature we dichotomized the mutations of each patient as clonal and subclonal, and we compared their contribution for each group (**Fig. 1D-J**). APOBEC mutational activity (SBS2 and SBS13) was typically enriched among subclonal mutations (**Fig. 1F**), while SBS9 was enriched among clonal mutations (**Fig. 1I**). This observation aligns with the MM pathogenetic model, which initiates in the GC with active SBS9 but not APOBEC (**Supplementary Fig. 4**)^43^. Subsequently, upon migration from a GC to the bone marrow, SBS9 becomes inactive, while APOBEC mutagenesis likely starts to cause mutations over time^60^. This temporal relationship was not seen in hyper-APOBEC patients, where SBS2 and SBS13 were enriched in both clonal and subclonal mutations, and SBS9 was often undetectable (**Supplementary Fig. 5**). This pattern supports a model where *MAF/MAFB* translocations induce a high level of APOBEC mutagenesis beginning in the early phases of tumor development. The melphalan signature, SBS99, was detected in both clonal and subclonal variants in RRMM patients. Interestingly, the majority of patients with SBS99 detectable in both clonal and subclonal variants were exposed to tandem HDM-ASCT (p=0.0069 compared to single HDM-ASCT using Fisher’s Exact Test), further confirming the cumulative mutagenic effect of melphalan on MM cells.

### Multiple myeloma molecular time

To investigate the temporal relationship between key genomic drivers in MM, we employed the recently published molecular time workflow^1,21,43,56^. This approach enables estimation of the relative timing of each chromosomal gain by using the corrected ratio between duplicated and non-duplicated mutations (**Fig. 2A-B; Supplementary Data 1**). For instance, if a chromosomal gain occurs early, few mutations are acquired before the duplication, resulting in a higher proportion of non-duplicated mutations largely acquired after the gain (purity-corrected variant allele frequency (VAF) ∼33%) compared to duplicated mutations (VAF ∼66%). Conversely, a similar proportion of duplicated and non-duplicated mutations is seen for late gains. A total of 332 (78.8%) samples from 314 (82.2%) patients qualified for the molecular time analysis, satisfying the criterion of having at least one large chromosomal gain (>1 Mb) with > 50 clonal SBS^21,56^. Among these, 249 (75%) and 83 (25%) were NDMM and RRMM, respectively. Molecular time data were generated for 1373 single gains (3:1, where 3 represents the total copy number and 1 the minor allele), 162 double gains (i.e., amplification, 4:1), and 147 copy neutral loss of heterozygosity (CN-LOH; 2:0), for a total of 1682 events (**Fig. 2C**).

**Figure 2:**
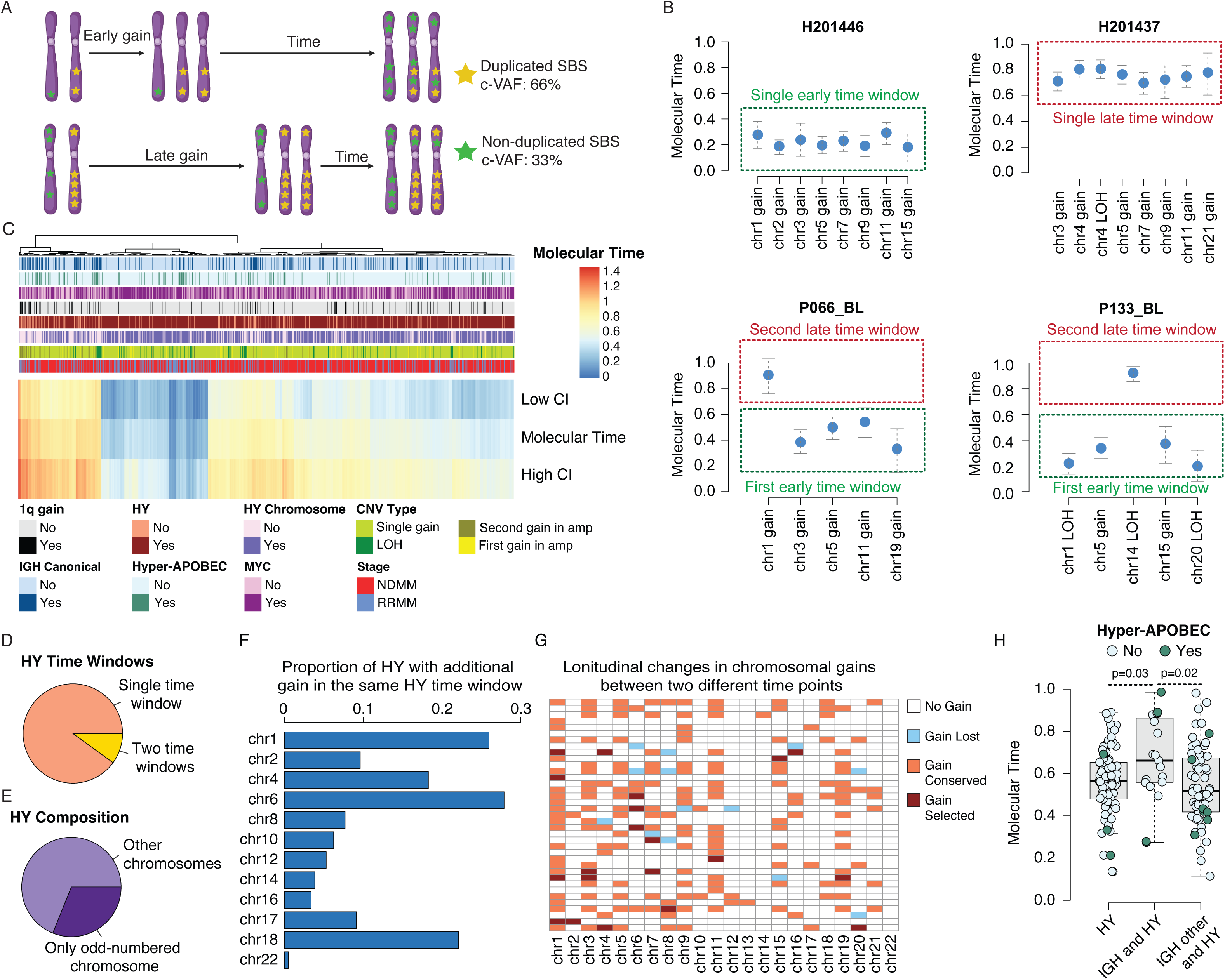
Molecular time and temporal relationship between large chromosomal gains in multiple myeloma. **A)** The cartoon summarizes the rationale behind molecular clock theory and how duplicated and non-duplicated mutations can be used to time the acquisition of evolutionary events. In each molecular time plot the blue dot represents the molecular time estimates, and the vertical line the CI generated by the *mol_time* bootstrap function. **B)** Example of different molecular time outputs in patients where multiple gains were acquired in the same time window or in two separate ones. **C)** Heatmap of all chromosomal gains (n=1682) with their molecular time and 95% confidence intervals (CI) across entire cohort. Each column represents a chromosomal gain for which molecular time data were generated. hyperdiploid (HY) chromosomes are defined as large gains involving 3, 5, 9, 11, 15, 19, and 21 (see **Supplementary Data 1**). **D-E)** Pie chart summarizing the proportion of HY samples where the final HY profile was acquired in two separate time windows **(C)** and those acquired simultaneously with other non-odd numbered HY chromosomes **(D)**. **F)** Bar chart showing the proportion of patients with non-HY chromosomal gains acquired in the same time window as the HY event. **G)** Heatmap showing the concordance of large chromosomal gains in 37 patients with more than one WGS sample collected at different time points (see **Supplementary Table 7**). Each row represent a patient. **H)** Boxplot of HY event molecular time calculated as the median molecular time of the earliest HY time window. Dark green dots represent gains belonging to samples that were hyper-APOBEC. Patients with hyper-APOBEC were usually grouped at the two extremes of the molecular time distribution because of their high mutational burden in either duplicated or non-duplicated variants. P-values were estimated using Wilcoxon test. IGH: canonical IGH translocations.

We initially examined the temporal relationship between chromosomal gains in 208 HY samples. If multiple chromosomal gains are detected in one sample, molecular timing can be used to estimate whether they were acquired within the same time window (similar molecular timing) or in two separate time windows (different molecular timing) (**Figure 2B**). In 91% (n=189) of the HY cohort, trisomies of odd-numbered chromosomes were acquired within the same time window (**Fig. 2C; Supplementary Table 5**). Of note, 62% of HY patients displayed additional chromosomal gains that were acquired within the same time window as the HY event, including gain of 1q (n=54) and 6p (n=58) (**Fig. 2D-F**). This finding suggests that the HY event is not limited to trisomies of HY chromosomes but may co-occur with other chromosomal segments. Furthermore, this data provides strong evidence that 1q gain can also be acquired in the very early phase of the disease. Among the 19 (9%) of HY patients in whom the odd-numbered chromosomal gains were observed in separate time windows, the most common was a late gain on chromosome 11 (n=10) (**Supplementary Table 5**). To validate the finding that HY patients can acquire gains of odd-numbered chromosomes in distinct time windows, we explored paired samples collected at different time points from 37 patients (**Fig. 2G, Supplementary Data 3** and **Supplementary Table 6**). Among the 21 HY patients, three exhibited at least one odd-numbered chromosomal gain acquired and/or lost at the second time point. In the complete set of 37 patients, chromosomal duplications that were either selected or lost exhibited significantly later molecular times compared to those that were shared between different time points, suggesting they were not present in the most recent common ancestor (MRCA) (see **Supplementary Figure 6A**). Furthermore, in patients where the final CNV profile was acquired through two temporally distinct windows, 88% (15 out of 17) of the unshared gains were acquired in the latest time window.

To further validate our fnding, we analyzed an independent FISH dataset of 75 patients with samples collected at the time of myeloma precursor conditions (i.e., MGUS and SMM) and NDMM. All patients were followed for at least five years at either the Mayo Clinic or the Heidelberg University, with at least two tests separated by at least five years. Overall, 3 out of 43 (7%) HY patients showed evidence of acquisition of additional odd-numbered chromosome gains over time (**Supplementary Table 7**).

Interestingly, we observed that patients with HY without canonical IGH translocations have a significantly earlier molecular time of the HY event compared to those with HY and concomitant canonical IGH translocations (p=0.02 using Wilcoxon test) (**Fig. 2H**). Both HY and non-HY gains detected in patients carrying a canonical IGH translocation were usually characterized by a later molecular time (p<0.001 for both using Wilcoxon, **Supplementary Fig. 6B-C**).

Of the three types of chromosomal aberrations studied (gain, amplification, CN-LOH), CN-LOH events accounted for 147 (9.1%) of them of which, 21 (14.3%) occurring in a second and later time window (where two independent time windows were present). A total of 96 CN-LOH occurred in 64 HY samples where we could estimate the molecular time of the HY multi-gain events. Among these, 23 (24%) affected the odd-numbered HY chromosomes (i.e., 3, 5, 7, 9, 11, 15, 19, 21). Intriguingly, 87 (27.9%) CN-LOH events were observed in the same time window as the HY multi-gain event in 27.8% of HY patients (n=58). 25.3% of these CN-LOH events were on odd-numbered chromosomes, reflecting a deletion of the non-duplicated allele. Importantly, CN-LOH on even-numbered chromosomes had a later molecular time compared with those on odd-numbered chromosomes in HY (p=0.0004 using Wilcoxon test; **Supplementary Fig. 6D**). Overall, this data indicates that CN-LOH, especially involving odd-numbered chromosomes often reflects a chromosomal duplication acquired within the context of HY multi-gain events. The deletion of the minor allele could be acquired either before or after the gain, however the current molecular time workflow does not allow for estimation of this temporal order of events. Finally, CN-LOH events were more frequently encountered in the RRMM samples, observed in 41% vs 18% of RRMM vs NDMM samples, respectively (p<0.001 using Fisher test) likely reflecting the impact of treatment in selecting late and subclonal events.

To interrogate these temporal patterns taking into account MM genomic heterogeneity, we investigated the molecular time patterns across the 12 genomically-defined MM biological subgroups (**Supplementary Fig. 1, Supplementary Table 4)** ^33^. Even after dividing patients with canonical IGH translocations or HY for the different patterns of genomic complexity, the *NSD2* [t(4;14)) and *CCND1* (t(11;14)] groups had an overall later acquisition of both all gains and odd-numbered HY gains compared to HY groups (**Supplementary Figure 7A-B; Supplementary Table 8**).

Next we focused our investigation on 1q gain, considering its increasing prevalence over the course of the disease and clinical prognostic relevance in MM. Overall, we observed an extensive heterogeneity across the 12 different biological subgroups, however no significant differences in the timing of 1q gain between these groups were observed (**Supplementary Figure 7C,** and **Supplementary Table 8**). Interestingly the chromosomal gains acquired within the three groups without canonical IGH translocations or HY [groups: Simple, Complex, and GainAmp1q_Del13q] were characterized by heterogenous temporal patterns, including early gains such as 1q. This suggests that 1q gain and non-HY genomic alterations may contribute to early disease pathogenesis, even in the absence of canonical IGH translocations and HY. The heterogeneous timeline of 1q gain also emerged from the FISH validation set, with 9 (12%) patients showing clonal 1q gain at both time points, and 13 (17%) patients showing 1q gain only at progression (**Supplementary Table 7**).

Overall, our temporal estimates suggest that canonical IGH translocations likely precede HY when both events are detectable eventually at time of diagnosis. They also suggest the existence of a distinct HY subgroup in which non-HY multi-gain events have an early, potential initiating role especially in the absence of canonical IGH translocations.

### Age of multiple myeloma initiation in patients’ life

To explore the temporal patterns of SBS signatures reported in **Fig. 1D**, we compared the mutational signatures landscape between duplicated and non-duplicated mutations (**Fig. 3**). To increase the accuracy of our SBS signatures estimates, only chromosomal gains with more than 50 mutations were considered^43,61^. SBS9 was particularly enriched in the pre-gain mutations within early duplications in HY patients. This pattern can only be explained with a model where HY is acquired in a B-cell already exposed to activation-induced deaminase (AID) and the light zone of the germinal center. SBS2 and SBS13 were found to be enriched in the post gain mutations in line with the late nature of this mutational process^43^. The only exception were patients with hyper-APOBEC where both signatures were equally detectable before and after the gains. Except for a minority of RRMM cases (n=7; 19%), SBS99 was usually only observed in post-gain mutations. SBS1 and SBS5 were detected in both groups of mutations, with more SBS5 in the non-duplicated vs the duplicated SBS. These patterns are consistent with the MM pathogenetic model propose in **Supplementary Fig. 4** based on SBS signatures differences between clonal and subclonal mutations.

**Figure 3:**
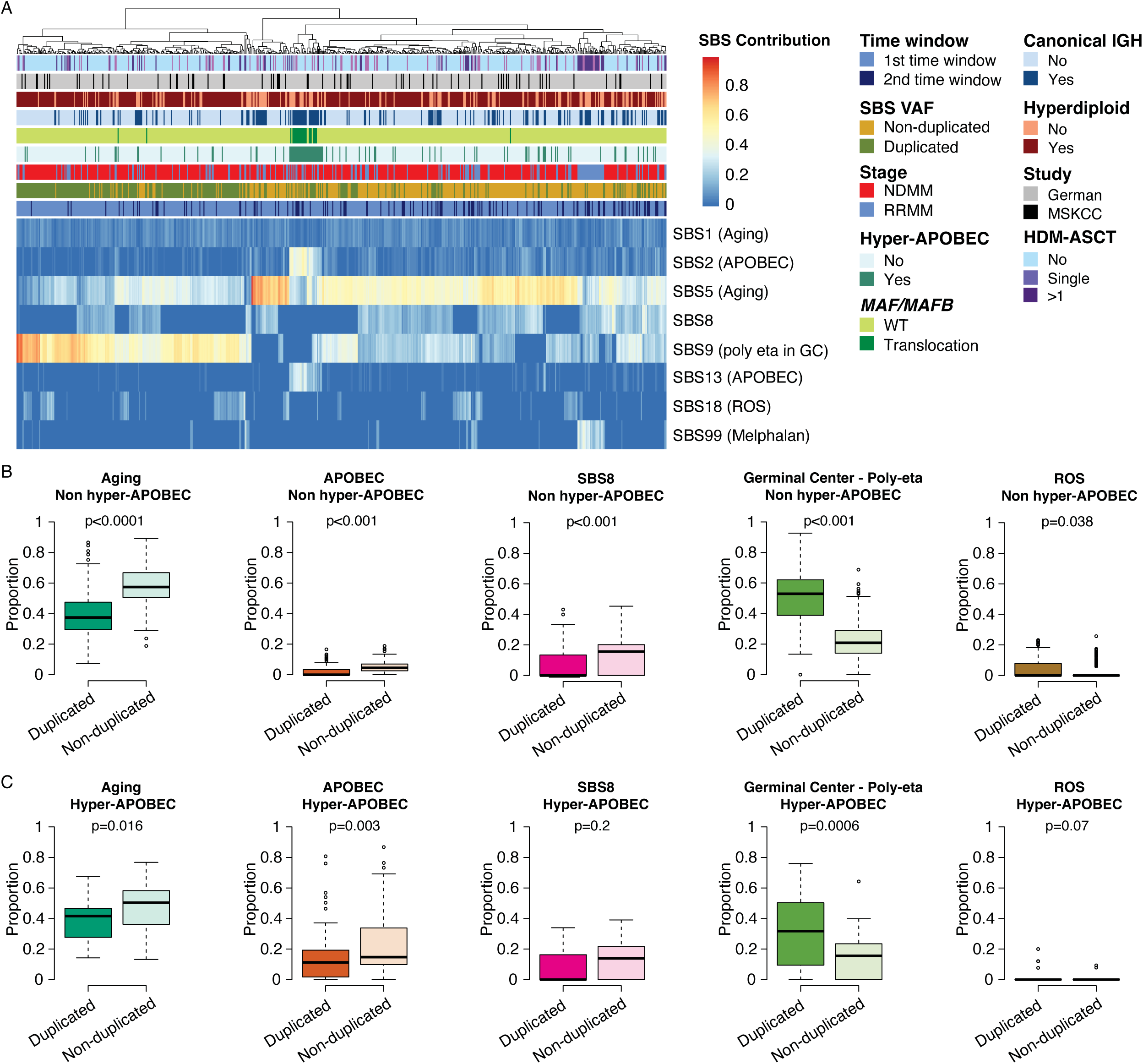
Timing multiple myeloma initiation. **A)** Heatmap summarizing the mutational signatures contribution among duplicated (i.e. pre-gain) and non-duplicated (i.e. post gain) mutations. SBS: single base substitution; GC: germinal center; ROS: radical oxidative stress; NDMM: newly diagnosed multiple myeloma; WT: wild type; RRMM: relapsed refractory multiple myeloma; HDM-ASCT: high dose melphalan and autologous stem cell transplantation; IGH: canonical IGH translocations; HY: hyperdiploid. **B-C)** Comparison of each signature’s contribution between duplicated and non-duplicated mutations. All non-hyper-APOBEC samples are plotted in **B)**, while **C)** shows the hyper-APOBEC samples. All p-values were estimated using Wilcoxon-test.

Above we showed that SBS1 and SBS5 clock like mutations are correlated with patients’ age in most of MM, B-cell lymhpomas and normal B-cells (**Supplementary Fig. 3**). Recently, the SBS1 and SBS5 molecular clock has been used in several tumors including a small MM series to estimate the absolute time in which chromosomal gains were acquired in a patient’s lifetime ^44,45,49,66^. To perform the same analysis in our large WGS series, we used the contribution for SBS1 and SBS5 before and after the gain to estimate the clock-like molecular time in each patient. To increase the accuracy of our mutational signature estimation, we first excluded patients in which their clock-like mutational burden did not fit in the linearity observed in the majority of patients (e.g. hyper-APOBEC; **Supplementary Figure 2B and Supplementary Data 2**). Secondly, we combined duplicated and non-duplicated mutations within chromosomes that were acquired within the same molecular time window. In line with the molecular time criteria, we only examined segments with a mutational burden greater than 50 in both the duplicated and non-duplicated mutations^43–45^. As previously shown ^21,44,45^, multiplying the clock-like molecular time of each time window for the patient age provided estimates of when these events were acquired in patient’s life (**Supplementary Data 1 and 2**). Our estimates among 155 NDMM patients who met these criteria showed that the median time of acquisition of the first set of gains in MM occurred in the second-third decades of life [median estimated age 26.5 years (range 7-62); **Figure 4A** and **Supplementary Table 9**]. The estimated median time lag between age at sample collection and the age in which the tumor cell acquired the first large chromosomal gain(s) was 32 years (range 1-57). Both the age of the first gain and the time lag were shorter in patients with canonical IGH translocation compared to HY without canonical IGH translocations (p<0.001, using Wilcoxon test);**Fig. 4B-C**). Overall, this suggests that HY is acquired later in the presence of pre-existing canonical IGH translocations, and that the interval between the late acquisition of HY and diagnosis may be shorter due to the more mature and temporally advanced genomic profile of these tumors.

**Figure 4:**
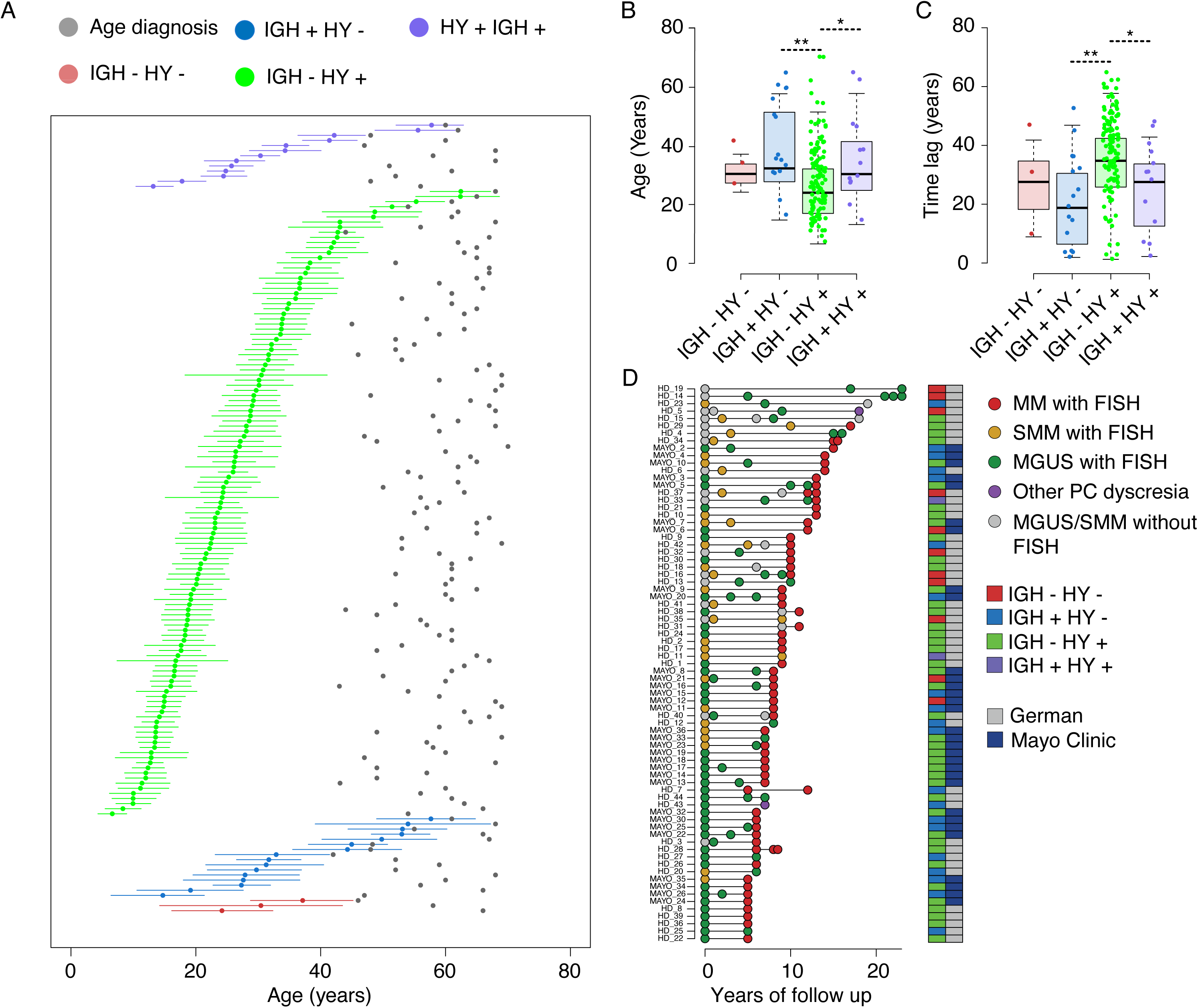
Timing multiple myeloma initiation. **A)** Absolute estimates and their confidence of interval (CI) of when the first large chromosomal gains were acquired in patients’ life. Estimates were generating calculating molecular time using SBS1 and SBS5 mutational burden among duplicated and non-duplicated mutations. SBS1-SBS5-based molecular time was calculated only in samples with >50 pre- and post-gain mutations. Only patients with residual <1900 in the linear model showed in Fig. 1B were included (n=155). **B-C)** Comparison of the estimated age of acquisition of the first gain (**B**) and the time lag between the age of diagnosis and the age at which patients acquired the first chromosomal gain (**C**) across different cytogenetic groups. * and ** reflect p-values estimated by the Wilcoxon test of <0.01 and <0.001, respectively. **D)** Longitudinal cohort of patients with samples evaluated by FISH for key genomic events, collected both at the time of myeloma precursor conditions and at diagnosis. The two FISH measurements were taken at least 5 years apart.

To further validate our estimates, we examined the independent FISH validation set (n=75, **Supplementary Table 7**). In all of these patients, canonical IGH translocations and/or HY were detectable at all time points examined, with 30 (40%) patients having these alterations in the precursor disease, 10-20 years before the MM diagnosis (**Figure 4D**). These findings, together with several previously published clinical evidences^1,5,6,19,20,27,28,41,67^, confirm that the evolution of MM extends over decades of a patient’s life.

### Temporal relationship between canonical IGH translocations and large chromosomal gains

Our initial analysis suggests that canonical IGH translocations precede HY in patients with both events. Yet, to provide more robust evidence for our interpretation, we identified 10 HY patients in whom a canonical IGH translocation was linked to a chromosomal gain. Among these, the most frequent occurrence was between t(11;14)(*CCND1*;*IGH*) and 11q gain, detected in 5 patients. Interestingly, two distinct temporal patterns were observed, depending on the type of translocation. In the first, 11q gain had an early molecular time and was directly caused by an unbalanced translocation between *CCND1* and *IGH*. In these cases, both events were acquired at the same time, likely at the beginning of tumor evolution (i.e., “unbalanced model”; **Fig.5A-B**). Conversely, in the second model, 11q gain had a late molecular time and was linked to a reciprocal translocation between *CCND1* and *IGH*. This scenario may be explained by the following temporal reconstruction: initially, a reciprocal translocation creates the t(11;14) and its derivative chromosomes. Subsequently, one of the two derivative chromosomes, particularly the one with 11q, is duplicated as part of a later event (i.e., “reciprocal model”; **Fig. 5C-D**). In both models HY gains were always acquired at a later molecular time compared to the t(11;14) supporting our initial interpretation. Expanding this concept to all patients with a t(11;14) and 11q gain irrespective of the HY status (n=19), gains on 11q acquired after a reciprocal translocation had a significantly later molecular time compared to gains acquired at the time of an unbalanced translocation (p<0.001 using Wilcoxon test; **Fig. 5E**). To validate the temporal order, we investigated an additional longitudinal FISH dataset from Mayo Clinic. This dataset includes 27 patients with t(11;14) and HY, tested at two time points: (i) MGUS/SMM and (ii) NDMM (**Supplementary Table 10**). The median time between the two samples was 3 years. In total, a t(11;14) was seen in all 27 patients as clonal at both stages. While four of these patients showed HY at both time points, two patients showed evidence of concomitant HY only at the MM stage, suggesting late acquisition of HY.

**Figure 5:**
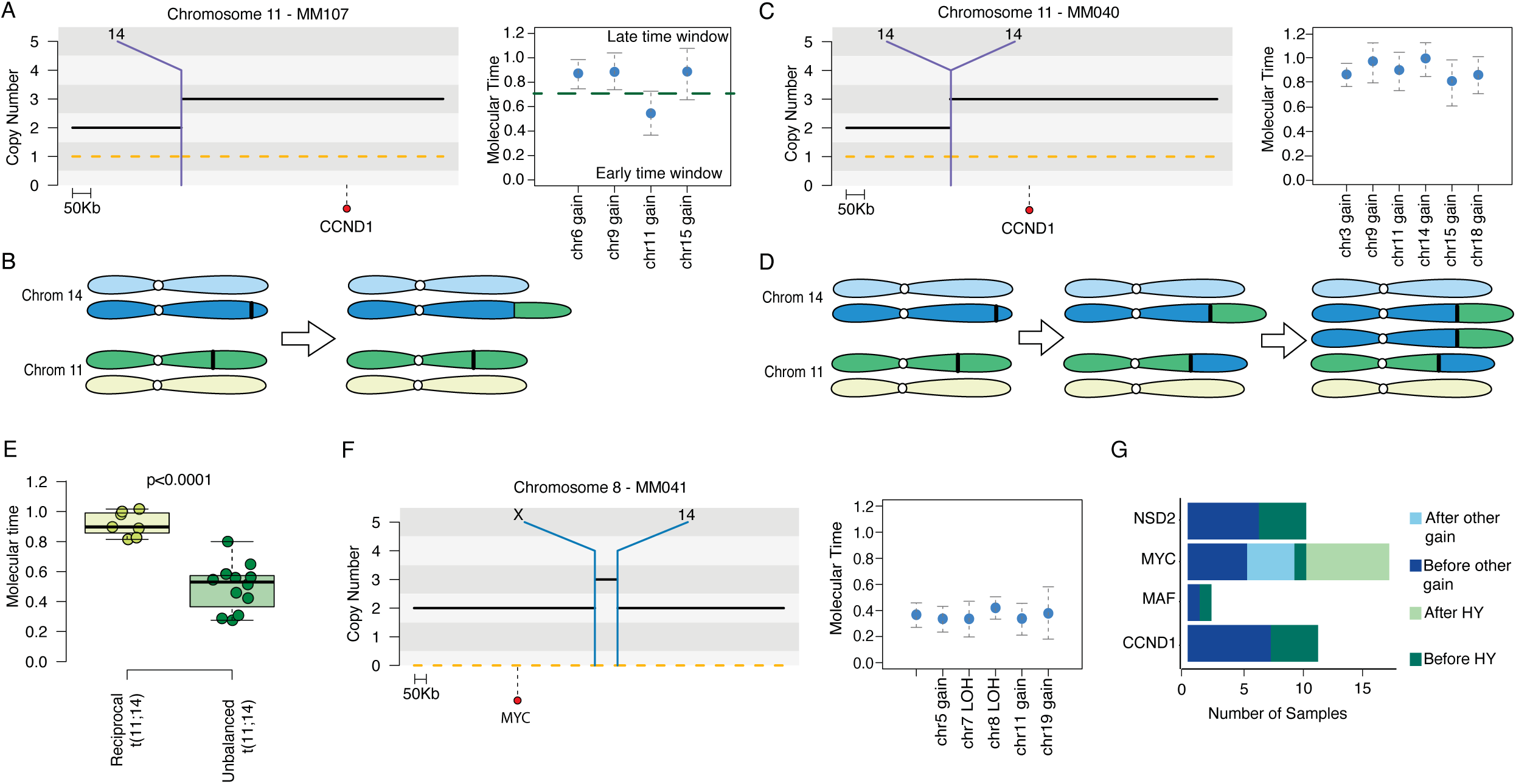
Timing of canonical IGH and MYC translocations. **A)** Unbalanced t(11;14)(*CCND1;IGH*) in a patient with hyperdiploid (HY) with zoomed-in view of breakpoint on chromosome 11. Molecular time of all chromosomal gains for this patient (right) shows earlier molecular time of chromosome 11 gain compared to other HY chromosomes (9 and 15). **B)** Cartoon depicting unbalanced t(11;14)(*CCND1;IGH*) occurring simultaneously and linked with chromosome 11q gain on derivative chromosome. Black lines represent translocation breakpoints. **C)** Reciprocal t(11;14)(*CCND1;IGH*) in a patient with HY with zoomed-in view of breakpoint on chromosome 11. Molecular time of all chromosomal gains to the right of the SV plot shows chromosome 11q gain with a similar molecular time as the other HY chromosomes (3, 9, and 15). **D)** Cartoon depicting reciprocal t(11;14)(*CCND1;IGH*) occurring as a multi-step process with translocation first followed by 11q gain on derivative chromosome as a separate and later event. **E)** Boxplot showing molecular time of chromosome 11q gain based on type of t(11;14)(*CCND1;IGH*) (reciprocal-left box; unbalanced-right box) with a later chromosome 11q gain with reciprocal translocation. P value was estimated using Wilcoxon test. **F)** Example of a patient with a templated insertion involving chromosomes 8, 14, and X on a copy neutral loss of heterozygosity (LOH) segment on chromosome 8, involving the *MYC* hotspot locus. Because the 3:2 CN jump, we can conclude that the CN-LOH preceded the templated insertion. The accompanying molecular time plot shows the CN-LOH involving chromosome 8 has a similar time to HY chromosomes in this patient, confirming the SV involving MYC is a late event. **G)** Horizontal bar plot depicting timing of canonical IGH translocations (*CCND1*, *MAF/MAFB*, and *NSD2*) and *MYC* SV relative to HY event and other chromosomal gains. In **A**, **C**, and **F** the black and orange dashed lines represent the total and minor copy number. Purple vertical lines represent translocations with the partner chromosome reported on the top of the line. In each molecular time plot the blue dot represents the molecular time estimates, and the vertical line the CI generated by the *mol_time* bootstrap function.

For the other IGH translocations, we identified only two cases (MM065 and MM064) where these two events could be linked with large chromosomal gains molecular time estimates. Using molecular time and recently published pre- and post-gain deletions workflow^68^ we demonstrated in both cases that IGH translocations preceed the gains (**Supplementary Fig. 8A-B** and **Supplementary Data 4**). The fact that IGH translocations are typically acquired before HY and other non-HY gains, and the fact that these gains are acquired more than 20 years before the diagnosis in patients with these translocations, imply that, similarly to HY, also canonical IGH translocations events can be acquired decades before MM diagnosis.

### Timing of MYC structural variants

Next, we investigated the temporal patterns associated with *MYC* SVs. In our cohort, we identified a total of 205 (48.7%) patients with such events, which were usually responsible for focal gains or deletions around the *MYC* locus. To time these events, we focused on patients in whom *MYC* SV-mediated focal gains occurred within a large chromosomal gain including 8q. In these patients, if the *MYC* focal gain caused a CNV jump of 1 (i.e. from 3:1 to 4:1), it meant that the *MYC* SV caused a focal gain on one of the two duplicated alleles, and was therefore acquired after the 8q gain (**Supplementary Fig. 8C-D**). In contrast, a CNV jump of 2 (i.e., from 3:1 to 5:1) meant that the tumor cell first acquired the SV-mediated gain and then duplicated the allele where the focal duplications were inserted (**Supplementary Fig. 8E-F**). This workflow was also applied to the partner chromosome of *MYC* translocations in case they had a focal gain within a large chromosome 8 gain. All 9 patients fulfilling the above criteria were HY. In 7 cases where chromosome 8 was gained in the same time window of HY, we showed that in 6 the *MYC* SV was acquired after HY (**Fig. 5F-G**). For example, in one patient (MM041), we observed a templated insertion involving *MYC*, chromosomes 14 and X. The focal gain near *MYC* was acquired on a large chromosome 8 CN-LOH with a CNV jump of 1 (i.e., from 2:0 to 3:0). This scenario can only be explained by a pattern where an LOH of this chromosome was acquired before the templated insertion (**Fig. 5G**). The HY karyotype occurred within the same time window as the LOH event on chromosome 8. Thus, we can conclude that the translocation involving *MYC* occurred after the HY initiating event.

### Chromosome 1q gain life history

Given its high prevalence and previously established prognostic impact^32,33^, we next investigated the temporal patterns of 1q gain. Overall, molecular time estimates were generated from a total of 88 and 25 patients with 1q gain and 1q amp (4 copies, 4:1 total alleles:minor allele), respectively. Cases with more than 4 copies of 1q were excluded as the molecular time could not be reliably estimated. The molecular time of both 1q gain and 1q amp was heterogeneous (**Fig. 6A-B**). In patients with canonical IGH translocations, 1q gain/amp was usually acquired late, while in patients with HY, 1q gain was detected either in the same time window (n=39) as the HY event or, as a late event in an independent time window (n=22). In 25 patients with 1q amp, the two gains were typically temporally unrelated, and in 14 (66%) patients, the first and second 1q extra copies were acquired in two clearly distinct and independent time windows. Together, these data further confirm that 1q gain can be acquired as either an early or late event during MM pathogenesis. Notably, the time of acquisition of 1q gain had a prognostic impact in our dataset. The molecular time of 1q gain was tested as a linear variable by cox proportional hazards and showed an association between early 1q gain and worse PFS and OS (p=0.03). Early 1q gain and 1q amp showed similar clinical outcomes (**Fig 6C-D**, where for graphical purposes, a cutoff of 0.85 was used to dichotomize groups; 1q early: ≤0.85, 1q late: >0.85). This difference in PFS was retained in a multivariate analysis including ISS stage and high-risk cytogenetic abnormalities (**Supplementary Tables 11-12**) ^32,33^. The only difference between early and late 1q gain was the presence of chromothripsis, which was higher in the early 1q gain group. However, this finding was no longer significant after correction for multiple testing (**Supplementary Table 13**). Overall, these data suggest that the adverse prognostic effect of numerical 1q aberrations in NDMM may depend more on the timing of their acquisition than on the number of extra copies gained.

**Figure 6:**
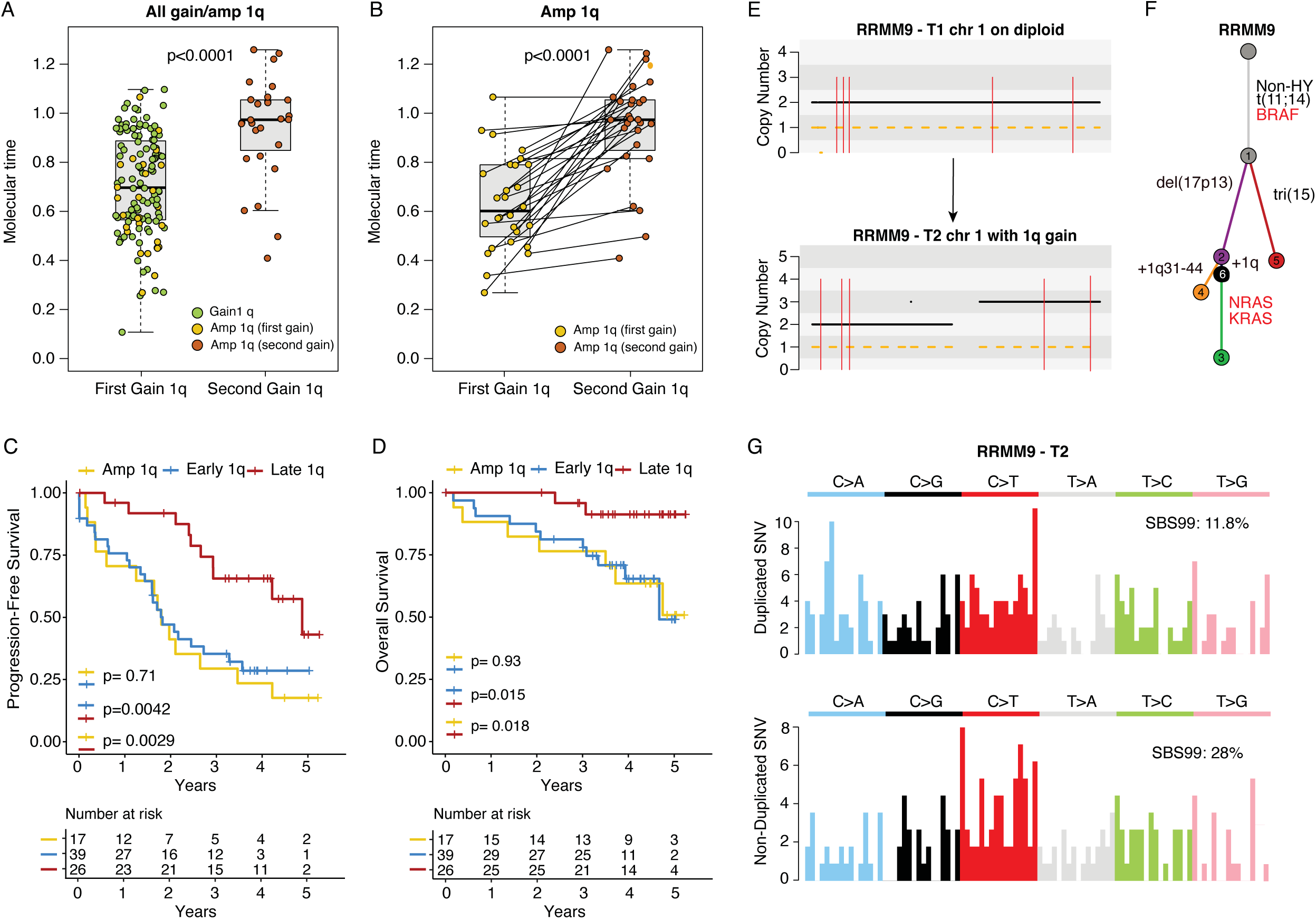
Timing and patterns of 1q gain and amplification. **A)** Boxplot of the molecular time of chromosome 1q gains divided into first gain (first box; single gains, or first gain of an amp) and second gain of an amp (second box) demonstrating a later molecular time for second 1q gains. **B)** Boxplot demonstrating the molecular time of first gain and second gain of 1q amp. Most paired gains had a significant difference in timing, with the second gain tending to occur at a later point in time compared to the first gain. In **A** and **B** p-values were estimated using Wilcoxon test. **C)** Kaplan Meier progression free survival (PFS) plot with 3 groups represented as follows: early 1q gain (molecular time <=0.85), late 1q gain (molecular time >0.85), and 1q amp. **D)** Kaplan Meier overall survival (OS) plot with the same 3 groups (early 1q gain, late 1q gain, 1q amp). In **C)** and **D)** he 0.85 was selected for graphical purpose. The association between early 1q and poot outcome was confirmed by coxph R function using molecular time linear feature. For 1q amplification, we exclusively considered cases with two extra copies and with molecular time estimates. **E)** Example of a relapsed refractory MM patient with a late 1q gain detected after exposure to high dose melphalan and HDM-ASCT, with both samples collected after HDM-ASCT. Top panel shows chromosome 1 as diploid at the earlier time point collected after HDM-ASCT. Our temporal estimates suggested that 1q gain was most likely already present, being acquired after HDM-ASCT. The bottom plot shows chromosome 1q gained at a second and later time point. Both samples were exposed to melphalan without any additional exposure between the two time points. The black and orange dashed lines represent the total and minor copy number. **F).** Phylogenetic tree for RRMM9 with key multiple myeloma annotated for each cluster. **G)** The 96 mutational classes for all chromosomes acquired in same time window as chromosome 1q gain for sample K08K-N5CC3E_T2 are reported on the bottom. Both duplicated (pre-gain) and non-duplicated (post-gain) show a pattern consistent with the SBS99 (melphalan) signature.

We next investigated whether 1q gain can be acquired during treatment as suggested by prior studies^69^. Therefore, we analyzed RRMM with 1q gain with evidence of the melphalan signature, SBS99, as consequence of HDM-ASCT. Since SBS99 is directly caused by melphalan, its acquisition by the tumor cell is restricted to a specific time window (i.e., exposure to melphalan before HDM-ASCT)^70^. If the 1q gain was pre-existing, then SBS99 would only be detected among the non-duplicated mutations on 1q. In contrast, if the 1q gain was acquired after HDM-ASCT, the SBS99 mutations on one allele would be duplicated and SBS99 would be detectable among the duplicated variants. Performing this workflow, we identified 3/26 RRMM patients where SBS99 was detected within duplicated variants, in line with acquisition of 1q gain after HDM-ASCT (**Fig. 6E-G**). Conversely, in the other 23 patients SBS99 was only detected within non-duplicated variants. Expanding the same analysis to other chromosomes, we found both gains (chromosomes 3, 6, 8, and 19) and CN-LOH events (chromosomes 9 and 16) that were acquired after exposure to melphalan. Overall, these data show that while the majority of numerical 1q aberrations are pre-existing, a fraction may be newly acquired during treatment, and the same is true for other chromosomal gains.

## DISCUSSION

Recently, the chronological order of genomic events in the molecular pathogenesis of MM and its precursors has been inferred from FISH and DNA sequencing studies. ^8,14,71^. From these studies, a number of patterns have been established, including 1) initiating events comprise canonical IGH translocations and HY; 2) numerical aberrations of 1q and *MYC* translocations are typically considered late-stage or events acquired/selected at progression; 3) 11q gain is always caused by t(11;14) when the two events co-occur ^2,3,20,22,28,29,31,37,55,72,73^.

In line with prior observations and to what seen in most of the cancers, we estimated that MM first events can be acquired 2-3 decades before the diagnosis. This mathematical estimation is supported by several epidemiological and clinical evidence across multiple tumors^44^, but in particular among MM. Indeed, it is noteworthy that there exists a precursor stage detectable several years before clinical manifestation in virtually all patients. In MM, evidence of monoclonal protein (i.e. SMM or MGUS) can be detected by serum protein electrophoresis more than 10-15 years before the diagnosis of MM. strongly supporting the evolution across a long time^5,6,38,67^. Furthermore, utilizing highly sensitive methodologies such as mass spectrometry, the Mayo Clinic group has demonstrated the detection of monoclonal protein up to 20 years prior to the diagnosis of MGUS and MM^41^. The long temporal evolution of multiple myeloma and the reliability of molecular clock estimates have also been validated using chemotherapy SBS signatures in patients who developed MM after being treated with platinum for ovarian cancer decades before^70^. To further validate the long MM time lag between initiation and diagnosis, we extracted a unique dataset of MM patients with available longitudinal FISH results at the time of diagnosis and during precursor conditions (i.e., MGUS and SMM). From this analysis, we confirmed that HY and canonical IGH translocations are usually conserved over time and can be detected 10-20 years before the diagnosis. Overall, these evidences support the idea that MM takes decades to develop.

Because MM evolution spans several decades and samples from very early and/or multiple time points are usually not available, several temporal and biological aspects of MM evolution remained unknown and awaited confirmation. To investigate these relevant aspects, we leveraged one of the largest WGS datasets for MM and timed individual chromosomal aberrations, enabling investigation into the temporal relationship between various key and highly prevalent genomic drivers in MM. Using this approach, we confirm prior assumptions and provide novel insight into the evolution of MM and prognostic impact of the temporal pattern of chromosomal changes. When applicable, our methods consistently confirmed that canonical IGH translocations, when present, likely represent the earliest event in the life history of MM. For the first time, we demonstrate that these translocations consistently precede large gains, including HY when they co-occur. Interestingly, HY molecular times were highly heterogeneous beyond the presence of canonical IGH translocations. This observation suggests that HY has a heterogenous role in MM pathogenesis, and may not always be an initiating event.

By studying the temporal relationship of odd-numbered chromosomal trisomies, we confirm that they are acquired simultaneously in the vast majority of HY patients. However, in ∼10% of patients, the final set of trisomies comprising the HY profile is acquired by temporally unrelated gains, highlighting the complex dynamisms that characterize MM life history^21^. We also show that LOH of odd-numbered chromosomes, as well as other chromosomal aberrations such as gain 1q or 6p, often co-occur with trisomies of odd-numbered chromosomes in HY, suggesting that a single catastrophic event may lead to multiple types of aberrations. These findings also suggest the importance of developing a comprehensive, genome-based definition of HY, that eventually includes LOH and gains of other chromosomes.

Recent longitudinal studies have identified branching evolution as the dominant pattern in patients who achieve deep responses to therapy ^21,37,59^. Yet, these studies have struggled to determine whether new lesions at relapse are due to selection of pre-existing occult subclones or acquisition of these lesions during treatment. Using SBS99 as a temporal barcode^74^, we were able to shed light on this crucial issue and show that ongoing genomic instability may indeed allow the acquisition of additional chromosomal aberrations, such as gain 1q.

The contention regarding numerical aberrations of 1q as a marker for high-risk MM has also been ongoing for more than a decade, and recent reports have even questioned its negative prognostic impact on outcome as an isolated feature^31,32,72,75–80^. In this study, we found that the prognostic impact of 1q appears to be more related to the time of acquisition rather than the number of copies. Considering the different temporal patterns, we observed that early 1q gain had a significantly worse outcome compared to late 1q gain. The adverse clinical outcome of early 1q gain was similar to that of 1q amp, where the first gain is usually also acquired early. This might explain the heterogeneity of the reported impact of 1q gains and amplifications, depending on the fraction of early gains in respective datasets^16,31–33^. Importantly, our dataset is composed of NDMM enrolled in three clinical trials and treated with current regimens, heightening the relevance of these results. Yet, we recognize that this finding is based on a limited sample set and will require validation in larger and independent cohorts.

In conclusion, in this study we have uncovered distinct temporal patterns of acquisition of genomic events that contribute to the reconstruction and better understanding of the life history of MM and may have potential clinical and prognostic implications.

## METHODS

### Patients

This study, adhering to the Declaration of Helsinki, received approval from Memorial Sloan Kettering Cancer Center and Heidelberg University. WGS from 421 samples were generated at MSKCC (n=60) and Heidelberg University (n=361).

### MSKCC cohort

MSKCC samples were collected within two phase II clinical trial (NCT03290950 and NCT02937571)^47,48,81^. The median coverage was 75X. The study size was decided based on samples availability. No statistical methods were used to pre-determine sample sizes. Frozen bone marrow cells from paired diagnosis and follow-up samples were thawed in warm Roswell Park Memorial Institute 1640 Medium (RPMI, Gibco, USA). Tumor cells from diagnostic samples were isolated from diagnostic samples by CD138 magnetic beads using either a QuadroMACS separator and LS column-based method (CD138 MicroBeads, human, Mylteni Biotec, USA) or a Robosep™ approach (EasySep™ Human CD138 Positive Selection Kit II, Stemcell, Canada) according to manufacturer’s instruction. Tumor cells from follow-up samples were sorted by flow cytometry based on the expression of CD38 (PE anti-human CD38 Antibody, Miltenyi, USA, 1:100), CD138 (APC anti-human CD138 Antibody, Biolegend, USA, 1:100), or CD45 (FITC Antibody, Miltenyi, USA, 1:200), with an additional Live/Dead exclusion (DAPI, Invitrogen, USA) using a BD FACS-Aria III cell sorter. FCR blocking was ensured using FcR Blocking (Human, Miltenyi). CD138-positive cells from diagnostic samples were enumerated on a Luna II cell counter and lysed in RLT Plus buffer (QIAGEN, Maryland, USA). The sample underwent homogenization (QIAshredder, QIAGEN), DNA extraction (AllPrep DNA/RNA Mini Kit) and quantification using both a photometric measurement (NanoDrop One/OneC instruments) and a fluorescent measurement (Qubit dsDNA BR Assay Kit or Qubit dsDNA HS Assay Kit, Thermofisher, USA). Additionally, matching peripheral blood mononuclear cells were lysed similarly, and DNA was extracted (QAmp Mini Kit); samples then proceeded for sequencing. Sixty matched tumor-normal samples underwent WGS at the MSKCC sequencing core. In short, following quantification via PicoGreen and quality control using an Agilent TapeStation, 100-1000 ng of genomic DNA was sheared (Covaris) and sequencing libraries were prepared using a modified KAPA Hyper Prep Kit (Kapa Biosystems). After ligation, libraries were subjected to a size selection using AMPure XP Beads (Beckman Coulter, #A63882). Libraries that required PCR amplification were pooled to the same concentration and non-PCR amplified libraries were pooled in the same volume. Samples were run on a NovaSeq 6000 in a 150 bp/150 bp paired-end run, using the NovaSeq 6000 SBS v1 kit and an S4 or S8 Flow Cell (Illumina). Target coverage depth was 80x for tumor and 40x for normal. Short insert paired-end reads were aligned to the reference genome (GRCh37) using the Burrows– Wheeler Aligner (v0.5.9). All samples were uniformly analyzed by the following bioinformatic tools^82^: somatic mutations were identified by integrating CaVEman, Mutect and Strelka. Only variants supported by at least 2 callers and not flagged by the other were included in the analysis. Copy number analysis and tumor purity (i.e., cancer cell fraction) were evaluated using ASCAT (https://github.com/VanLoo-lab/ascat) and Battenberg (https://github.com/Wedge-Oxford/battenberg). SV were defined by BRASS (https://github.com/cancerit/BRASS) via discordant mapping of paired-end reads^26^.

### Heidelberg University cohort

Among the 361 WGS enrolled from Heidelberg University 256 NDMM samples as well as the corresponding germline control (mainly peripheral blood) came from the GMMG-HD6 clinical trial^46^. Mononuclear cells from bone marrow aspirates as well as from peripheral blood were isolated using the Ficoll-Pague method. Bone marrow myeloma cells were enriched using human CD138 Microbeads and Manual MACS Cell Separation Units (Miltenyi Biotec, Bergisch, Gladbach, Germany). DNA of CD138-positive fractions was isolated using the Allprep Kit (Qiagen). WGS libraries were prepared with the Illumina TruSeq Nano DNA kit and sequenced on Illumina NovaSeq 6000 Paired-end 150 bp S4 to an average coverage of 88x for tumor and 47x for germline control samples, respectively. Raw sequencing data was processed and aligned to human reference genome build 37 version hs37d5 using the DKFZ OTP WGS pipeline.^83^ CNV, estimation of tumor ploidy and purity were determined with ASCAT. Indels were called by Platyphus^84^. SBSs were determined using samtools mpileup (v1.2.166-3)^85^. For the indels and SBSs additional filtering steps were applied, including blacklist filtering,^84^ fpfilter (https://github.com/genome/fpfilter-tool, only SBS) and removal of events located in regions coding for immunoglobulins. SV were called using SOPHIA (https://github.com/DKFZ-ODCF/SophiaWorkflow).

### Construction of phylogenetic trees

For the construction of phylogenetic trees, mutations of paired samples were first analyzed using the R package MOBSTER v.1.0.0 to discriminate positive subclonal selection from neutral tails ^86^. However, mutations that were classified as tail mutation in one sample, but assigned to cluster C1 in the paired sample were not removed. Clusters were determined using the R package VIBER v.0.1.0, and clonal ordering and visualization were performed with the R package clonevol v. 0.99.11.

### Mutational Signatures

Mutational signatures were comprehensively analyzed across all whole genomes using a three-step process involving de novo extraction, assignment, and fitting ^50,87^. In the initial step, *SigProfiler* (https://github.com/AlexandrovLab/SigProfilerExtractor) was utilized for the extraction of SBS signatures^52^. These extracted signatures were then cross-referenced with the latest Catalogue of Somatic Mutations in Cancer (COSMIC) (https://cancer.sanger.ac.uk/cosmic/signatures/SBS) to identify known mutational processes active within the cohort. Subsequently, we employed *mmsig* (https://github.com/UM-Myeloma-Genomics/mmsig) as a fitting algorithm in the final step^88^. This algorithm confirmed the presence and estimated the contribution of each mutational signature in every sample, guided by the catalog of signatures extracted for each respective sample. The SBS-MM1 melphalan signature, which was reported previously, is now incorporated into the COSMIC catalogue of SBS signatures as SBS99.

### Molecular time

The temporal sequence of copy number and structural variant events was assessed using the R package mol_time (https://github.com/UM-Myeloma-Genomics/mol_time) ^35^. This approach enabled the estimation of the relative timing of large chromosomal gains (>1Mb), clonal chromosomal gains (3;1 or 4:1), trisomies (3:1), LOH (2:0), and amplification (4:1). The estimation relied on the purity-corrected ratio between duplicated clonal mutations. Clonal SBS were categorized into duplicated or non-duplicated based on the VAF corrected for cancer purity. The latter was estimated by combining purity estimates from both ASCAT and SBS VAF density and distribution within clonal diploid regions. Segments with more than 50 clonal mutations, excluding immunoglobulin loci and kataegis events, were considered for copy number alteration (CNA) analysis. Tetrasomies (2:2) were excluded due to the challenge of determining whether both chromosomal gains occurred in close temporal succession. The relative molecular time of each gained CNA segment was estimated using the described approach, distinguishing between chromosomal gains occurring in the same or different time windows. To determine the absolute time of acquisition for large chromosomal gains, we pooled pre- and post-gain mutations from chromosomal gains acquired in the earliest time window for each patient. Subsequently, we estimated the contribution of SBS1 and SBS5 using *mmsig* for both groups. Only set of gains with more than 50 mutations were considered. By leveraging the clock-like mutational burden of SBS1 and SBS5 among duplicated and non-duplicated mutations, we recalculated the molecular time. Molecular time based on SBS1-SBS5 was calculated only for samples with more than 50 pre- and post-gain SBS1-SBS5 mutations. Multiplying this clock-like molecular time by the age of sample collection provided estimates for when the gain was acquired ^43,45,56^. Samples with a residual >1900 in the linear model, considering age, SBS1-SBS5 mutational burden corrected for stage and hyper-APOBEC (**Fig. 1B**), were excluded from the absolute timing analysis.

To estimate the timing of loss-of-function events and the acquisition of distinct SV (e.g. IGH translocations), two approaches were employed ^89^: 1) linking SV breakpoints to the molecular time of chromosomal gains caused by the same SV; 2) estimating the relative time of SVs within large chromosomal gains based on the copy number of the SV breakpoint.

### Retrospective review of Fluorescence in situ hybridization (FISH) data

Interphase FISH analysis at Mayo clinic was performed on bone marrow cells using standard fluorescence *in situ* pretreatment, hybridization, and fluorescence microscopy ^90,91^. The ploidy of malignant PCs was assessed by either FISH or by flow-cytometry-based assessment of DNA index^92^. Review of the historical FISH database (2004-2024) including Mayo clinic patients with multiple FISH testing instances revealed approximately 10,000 records. Cases with an abnormal FISH result prior to MM diagnosis and at the time of MM diagnosis were identified. Those with evidence of t(11;14) positivity at the time of FISH testing were selected, included those cases with an abnormal FISH result >5 years prior to MM diagnosis (regardless of the FISH abnormality) were also selected and analyzed. This analysis was approved by Institutional Review Board from the Mayo Clinic (17-000820).

Interphase FISH analysis at Heidelberg University was accomplished on CD138-purified plasma cells as described previously^93^ using probes for the detection of numerical aberrations of the chromosome regions 1q21, 5p15/5q35 (only if necessary to define hyperdiploidy), 6q21, 8p21, 9q34, 11q23, 13q14.3, 15q22, 17p13, 19q13, and 22q11, as well as for the IgH translocations t(11;14)(q13;q32), t(4;14)(p16.3;q32), and t(14;16)(q32;q23). The score of Wuilleme et al was used to assess ploidy ^79^. Gains of at least 2 of the 3 chromosomes 5, 9, and 15 were used for a FISH definition of hyperdiploidy.

### Statistical analysis and plotting

Proportional testing was performed using Wilcoxon and Fisher’s exact tests to compare the median of a continuous variable or the distribution of discrete variables across groups, when appropriate. All p-values are two-sided if not specified otherwise. For hierarchical clustering, Euclidean distance was used as the default distance measure within the pheatmap R package. The Kaplan–Meier estimator was used to calculate time-to-event distributions. PFS was measured from the date of start of treatment to the date of progression or death, whichever occurred first. All plots and figures were generated using R-Studio Version 2023.09.1+494 (2023.09.1+494), Adobe Illustrator, and Biorender.

## DATA SHARING STATEMENT

GMMG-HD6: are currently in the process to be uploaded on EGA: EGAS00001007469 MSKCC WGS data have been uploaded on EGA: EGAD00001011132. RRMM WGS data have been uploaded on EGA: EGAS00001006538, EGAS00001004363, EGAS00001004805 and EGAS00001005973.

## AUTHOR CONTRIBUTIONS

F.M., M.S.R., and N.W., designed and supervised the study, collected, generated, and analyzed the data and wrote the paper. M.K. and A.M.P. collected, generated, analyzed the data, and wrote the paper. B.Z., A.C., analyzed the data. N.K., S.U., O.L. E.K.M., H.G., K.C.W., A.L., R.F. supervised the clinical trials, collected data. V.R., S.K., L.B. collected and analyzed the FISH data from Mayo Clinic. K.M., M.C., B.D., P.B., L.J., P.R., .S.H., M.A.B., D.G., Y.Z., .F.D., G.M., A.D. collected the data.

## ACKNOWLEDGMENTS

This work was supported by the Myeloma Solutions Fund (MSF), Paula and Rodger Riney Multiple Myeloma Research Program Fund, the Tow Foundation, Sylvester Comprehensive Cancer Center NCI Core Grant (P30 CA 240139), Memorial Sloan Kettering Cancer Center National Cancer Institute (NCI) Core Grant (P30 CA 008748), and NYU NCI Core Grant (P30CA016087). FM is supported by Leukemia & Lymphoma Society and by International Myeloma Society (IMS). GJM received grant support through a Translational Research Program award from the Leukemia & Lymphoma Society (6020-20). KM received grant support from the Royal Australasian College of Physicians, the American Society of Hematology and the Multiple Myeloma Research Foundation. AMP is funded by the Medical Data Scientist Programm of Heidelberg University, Faculty of Medicine. The Heidelberg Team thanks the Sample Processing Lab (SPL), the High Throughput Sequencing unit of the Genomics & Proteomics Core Facility and the Omics IT and Data Management Core Facility (ODCF) of the German Cancer Research Center (DKFZ), the DKFZ-Heidelberg Center for Personalized Oncology (DKFZ-HIPO) office, the Biobank Multiple Myeloma UKHD and the Myeloma Registry for excellent services. Support and funding of the project via the Dietmar-Hopp Foundation and the NCT Heidelberg Molecular Precision Oncology Program (project K08K) is gratefully acknowledged. Data storage service via SDS@hd is supported by the Ministry of Science, Research and the Arts Baden-Württemberg (MWK) and the German Research Foundation (DFG) through grants INST 35/1314-1 FUGG and INST 35/1503-1 FUGG.

