## Supplementary Data 1 for "Temporal Genomic Dynamics Shape Clinical Trajectory in Multiple Myeloma"

### Supplementary Data 1 – Timing multiple myeloma evolution

#### *Introduction.*

Multiple myeloma (MM) is the second most common hematological cancer, characterized by the aberrant proliferation of clonal plasma cells in the bone marrow, leading to organ damage such as lytic lesions, kidney failure, and bone marrow failure. Since the 1980s, it has been established that MM is consistently preceded by the asymptomatic expansion of clonal plasma cells, termed either monoclonal gammopathy of undetermined significance (MGUS) or smoldering myeloma (SMM). While these myeloma precursor conditions (MPC) are found in 3-5% of the adult population, only a small fraction will ultimately progress to MM and require treatment. Historically, using serum protein electrophoresis and free light chains, monoclonal protein could be detected decades before diagnosis. Recently, novel approaches, such as mass spectrometry and clinical trials, have shown that the monoclonal protein can be detected 1-2 decades before MGUS diagnosis, further supporting a long evolutionary period. The progression from germinal center initiation to multiple myeloma is usually characterized by an increase in the monoclonal protein produced by the clonal aberrant plasma cells. This increase can follow different models, but it is generally characterized by a multi-step evolution with phases of expansion and stability (**Supplementary Data Figure 1**). This behavior is interpreted as the result of the progressive acquisition and selection of distinct genomic drivers. Various cytogenetic, gene expression, and next-generation sequencing assays have established that the genomic evolution of MM is characterized by the accumulation of different drivers over time. Some events are defined as early genomic defining events (e.g., IGH translocations and hyperdiploidy), while few others are enriched in MM or in SMM with imminent risk of progression (e.g., MYC, MAPK mutations, and TP53). Despite validation in multiple independent studies, the temporal dynamics of most myeloma genomic defining events remain poorly defined.

**Supplementary Data Figure 1. Cartoon summarizing the pathogenetic role**

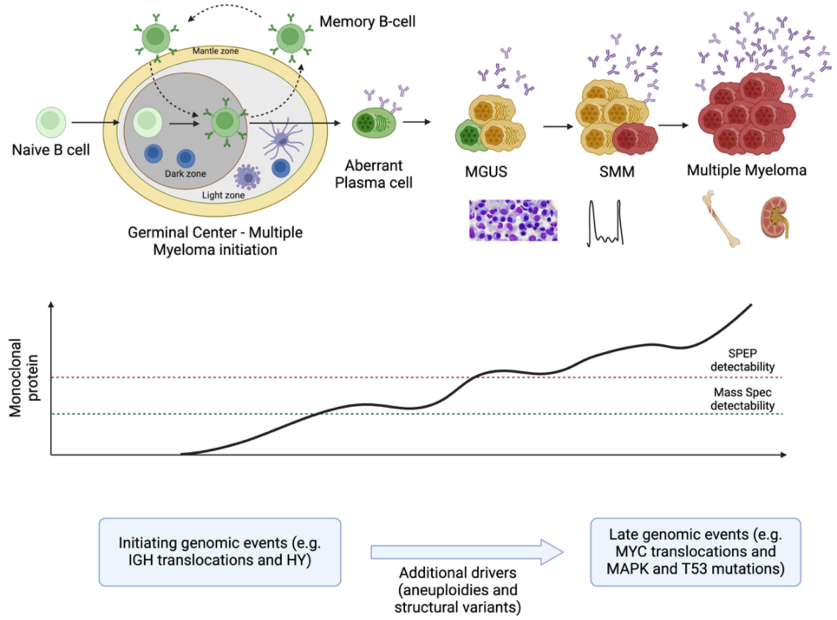

#### *Multiple myeloma biological subgroups: hyperdiploid and IGH translocations.*

MM has been historically divided into two main biological subgroups defined by distinct genomic events: hyperdiploid (HY) and those with IGH translocations. HY, detectable in 60-70% of newly diagnosed multiple myeloma (NDMM), is characterized by the duplication of multiple odd-numbered chromosomes (e.g., trisomies involving at least two chromosomes among 3, 5, 7, 9, 11, 15, 19, and 21; see **Supplementary Data Figure 2**). IGH translocations are usually detectable in 40% of NDMM cases and cause upregulation of key oncogenes by hijacking immunoglobulin regulatory regions and juxtaposing them next to these genes. The most important and frequent IGH translocations involve *CCND1*, *NSD2*, *MAF*, and *MAFB*. Based on their impact on gene expression, their persistent clonality over time, and their detection in myeloma precursor conditions (MGUS and SMM), these events have been proposed as “initiating events” in myelomagenesis. While this assumption is likely true for most patients, ~10% of NDMM cases can have both IGH translocations and HY. Additionally, other non-odd-numbered chromosomes can be duplicated and remain persistently clonal over time. These observations suggest that early myelomagenesis and the life history of myeloma may be more complex than previously thought. In fact, in a patient where both HY and 1q gain are clonal, it is not possible to use the copy number change cancer cell fraction to establish the order of event. 1q gain could have preceded the HY, or HY occurred earlier, or both events occurred at the same time.

**Supplementary Data Figure 2. Examples of copy number profile of multiple myeloma patients with available WGS.**

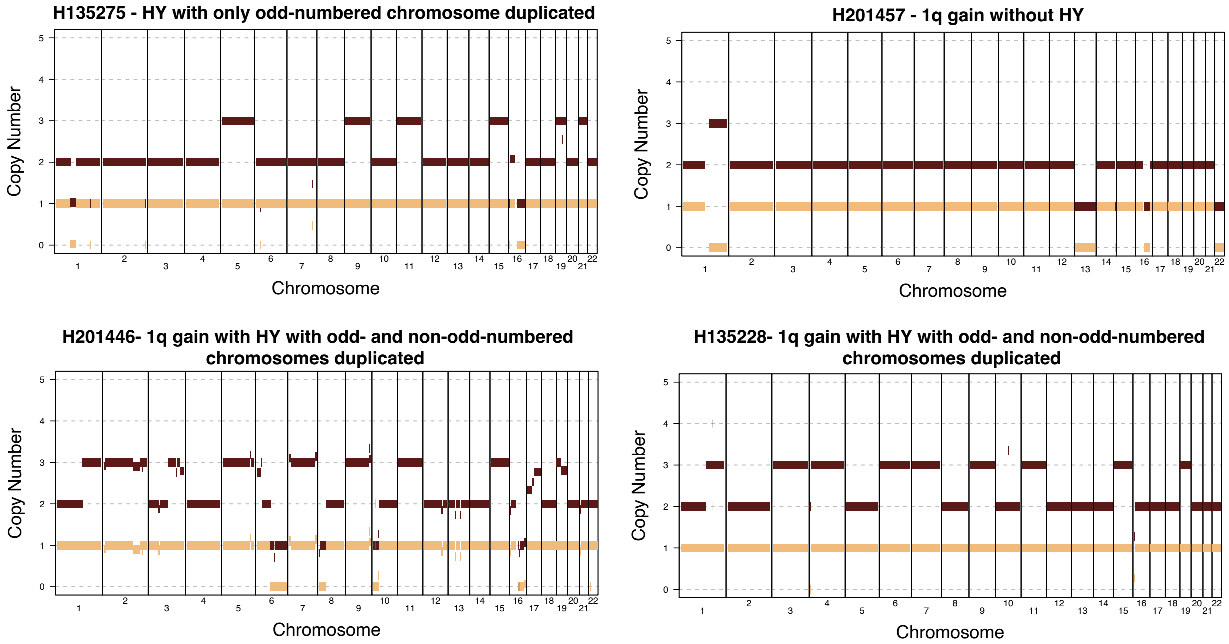


#### *Molecular time*

Tumors contain a record of all mutations acquired throughout their life history, which can be reconstructed from whole genome sequencing (WGS) data into a phylogenetic tree. The trunk of this tree encompasses all mutations acquired from the fertilized egg through transformative events up to and including the most recent common ancestor (MRCA). From the MRCA, the evolutionary trajectory diverges into one or more distinct branches or subclones, detectable by

WGS. The timing of landmark events along the phylogenetic tree can be understood in terms of molecular and chronological time.

Molecular time refers to the relative order of events along the cancer phylogenetic tree, estimated here by the corrected ratio of duplicated and non-duplicated mutations within large clonal chromosomal gains. The differentiation between duplicated and non-duplicated mutations is based on the variant allele frequency corrected for purity (c-VAF). If a mutation is acquired before the gain on the allele that will be duplicated, the VAF at the time of WGS and after the duplication will be 66%. In contrast, mutations acquired after the gain in one of the two duplicated alleles or at any time in the minor allele will have a c-VAF of 33%. The molecular time workflow utilizes the ratio of duplicated to non-duplicated mutations to gauge the temporal sequence of these genetic alterations within a patient's life. With individual molecular time estimates assigned to each chromosome, comparisons can be made across a spectrum of chromosomal gains within the same patient, allowing for the determination of temporal proximity or lack thereof between these events. This workflow, widely used in recent studies of both tumor and normal samples, indicates that a gain occurring early in time will have a low duplicated mutation burden (see **Supplementary Data Figure 3**). To ensure the accuracy of the model, we tested only clonal chromosomal gains larger than 1 Mb with more than 50 clonal mutations.

**Supplementary Data Figure 3. Rational behind molecular time workflow for single gain.** Next to the plot we reported the mathematical formula used for single gain.

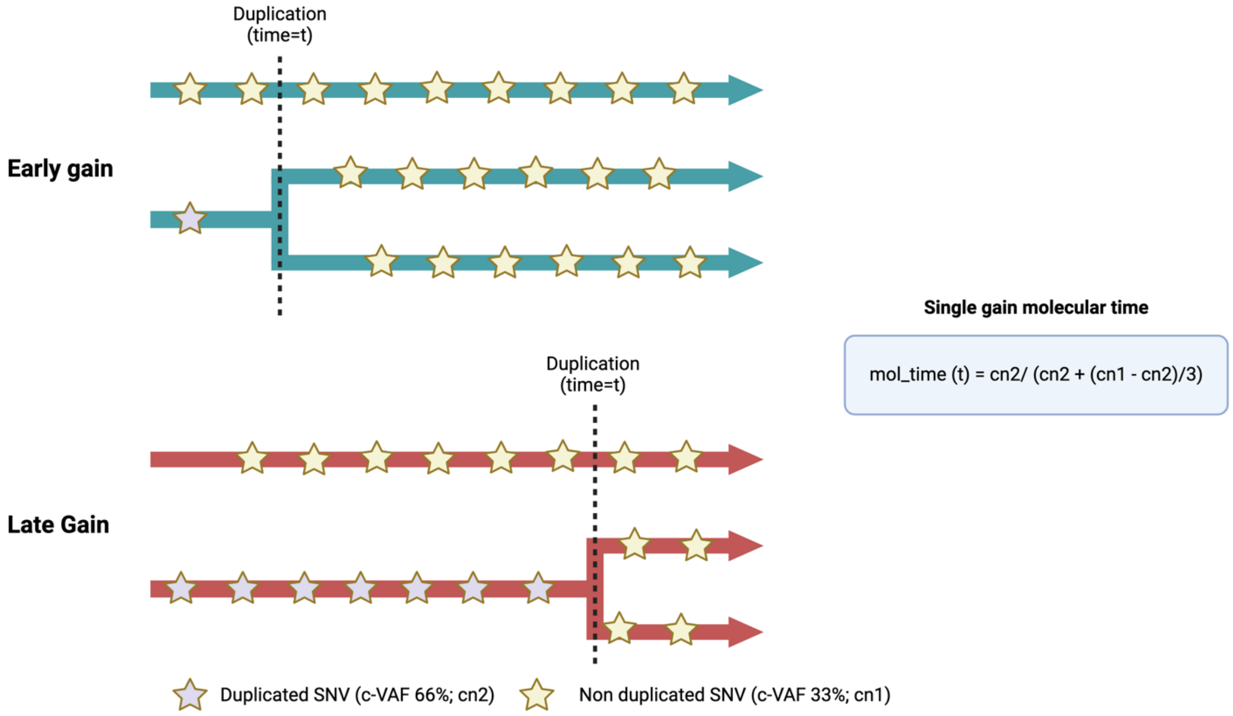


The model can also be extended to analyze copy-neutral loss of heterozygosity (LOH) and amplifications, such 4;1, where 4 is the total number of alleles and 1 is the minor (refer to **Supplementary Data Figure 4**). In these instances, the model will provide two molecular time estimates for each duplication event. It's worth noting that tetrasomies, where both parental alleles were duplicated, were excluded from consideration due to the inability to ascertain whether the duplications occurred simultaneously or not.

**Supplementary Data Figure 4. Rational behind molecular time workflow for amplifications (4;1) and LOH (2:0).** Next to the plot we reported the mathematical formula used for single gain.

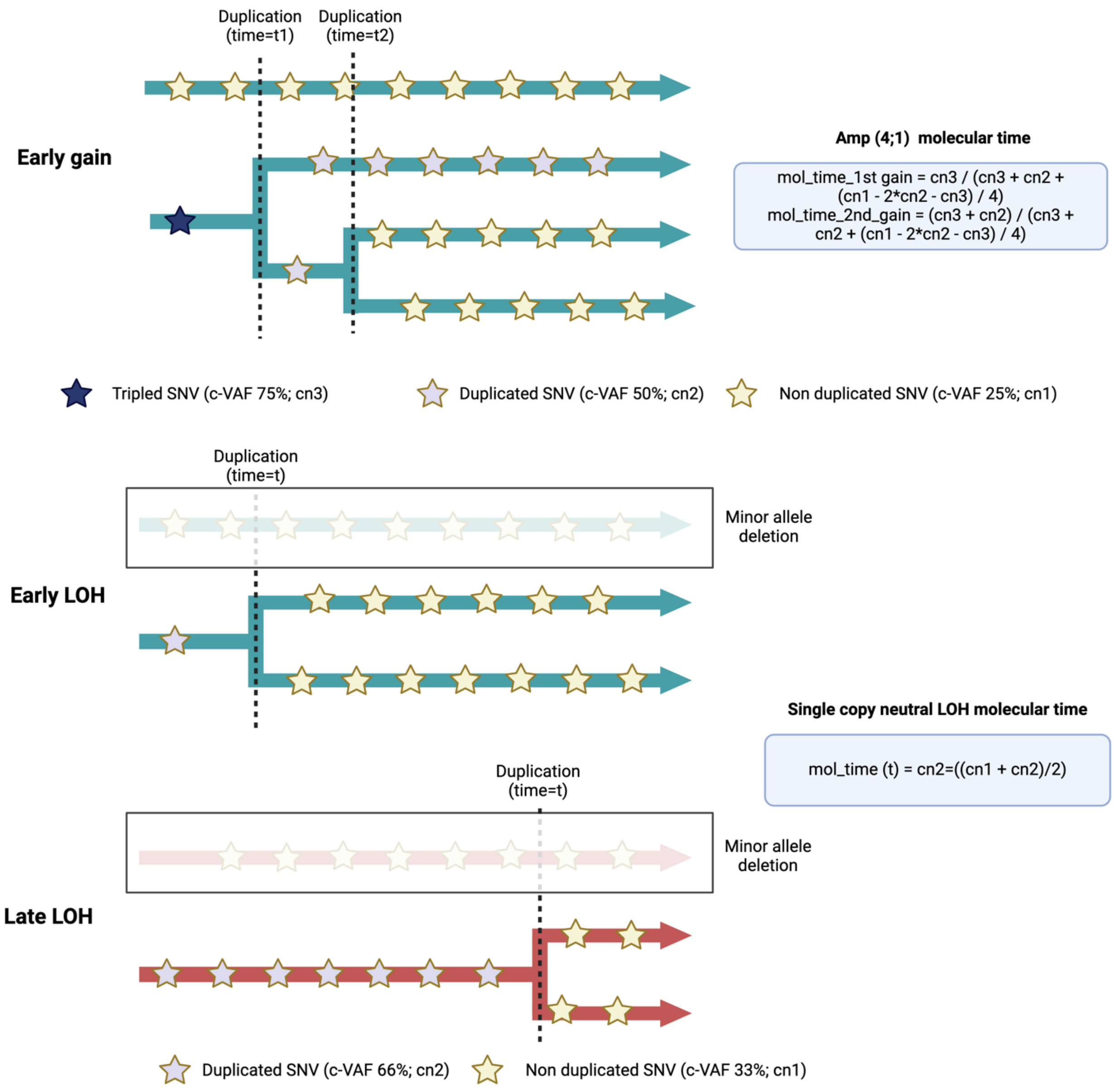


Employing the molecular time workflow in patients with HY enables the establishment of the chronological sequence in which large chromosomal gains occurred. Through this process, we employed a bootstrap function to assess whether different gains occurred in close temporal proximity. Gains deemed close in time were grouped within the same time window, while those outside of it formed new time windows. For instance, in the case of a patient with HY, molecular time analysis revealed that all duplicated chromosomes larger than 1 Mb with over 50 clonal mutations experienced gains that were closely aligned in time. However, it's important to note that molecular time analysis cannot definitively determine if these gains originated from a single mitotic event. Importantly, the molecular time approach offers a reliable and reproducible methodology, underscored by the clear differentiation between duplicated and non-duplicated corrected variant allele frequencies (VAF), as demonstrated in **Supplementary Data Figure 5**.

**Supplementary Data Figure 5. Example of molecular time analysis in hyperdiploid multiple myeloma. A)** Copy number profile of a newly diagnosed multiple myeloma patient. **B)** Molecular time estimates for each large chromosome with more than 50 clonal mutations. Each blue dot represents a molecular time estimate, with the confidence interval reflecting the bootstrap results from the molecular time function. **C)** Distribution of duplicated (green) and non-duplicated (red) mutations across each duplicated chromosome with a molecular time estimate.

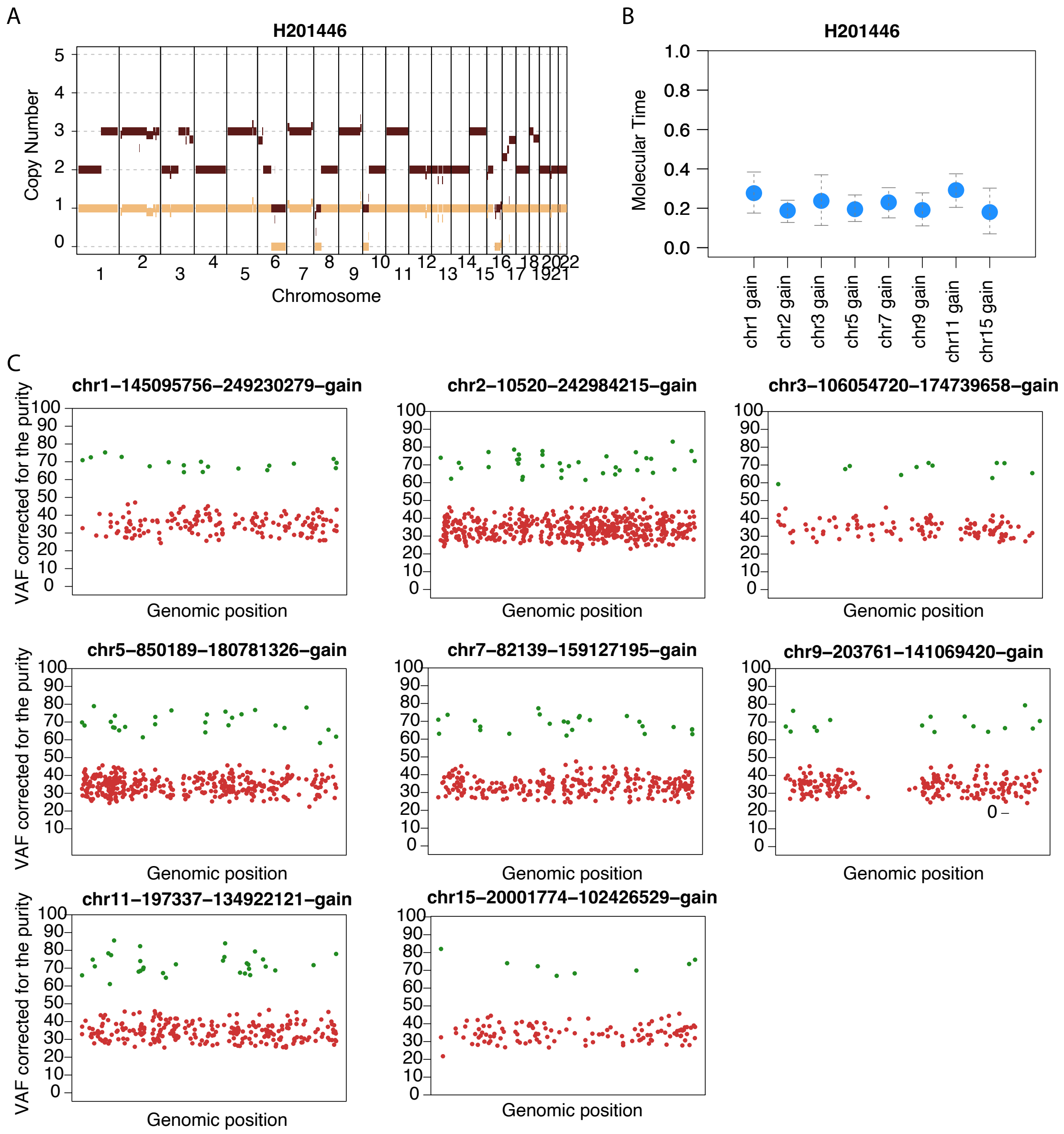


There are instances where not all odd-numbered chromosomes are acquired within the same time window or HY event. In the example below (**Supplementary Data Figure 6**), chromosomes 7 and 3 exhibit significantly different burdens of duplicated mutations. Consequently, the molecular time for chromosome 7 is higher, indicating a later acquisition independent of the multi-gain events during which chromosome 3 was acquired.

**Supplementary Data Figure 6.** The plot on the left shows the distribution of duplicated and non-duplicated mutations across chromosome 3 and 7. Next the molecular time output, where the dashed green line divides the two independent time windows.

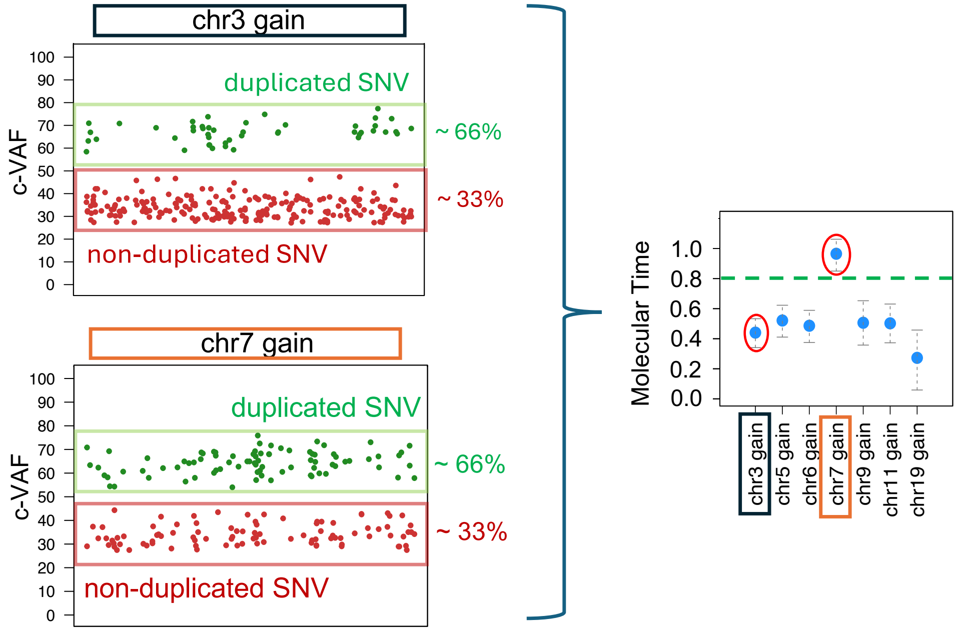


We can extend the application of molecular time analysis to non-odd-numbered chromosomes as well. In the example below (**Supplementary Data Figure 7**), we examine a hyperdiploid (HY) patient with a 1q gain. Given that all chromosomes are clonal, the final copy number variation (CNV) profile could result from three different evolutionary paths: 1) the 1q gain occurred after the HY event; 2) the 1q gain occurred before the HY event; or 3) the 1q gain and the HY event occurred simultaneously. Using molecular time analysis, we demonstrated that the 1q gain and the HY event occurred within the same time window, suggesting they were acquired early and in close temporal proximity.

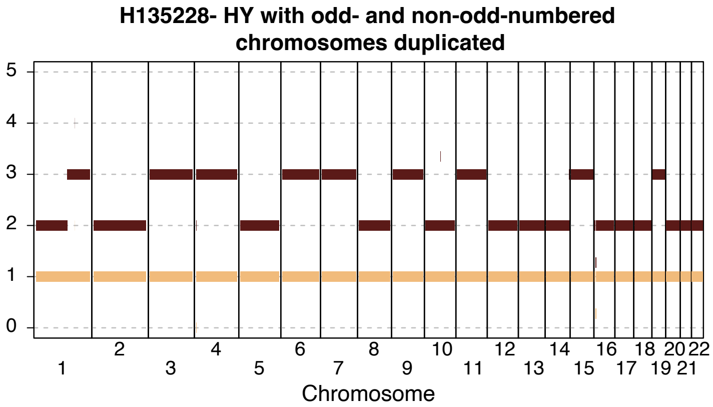


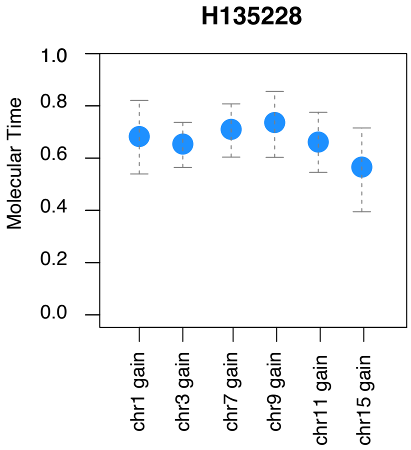


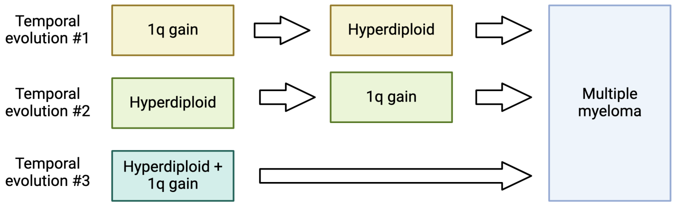


Molecular time shows that Temporal evolution #3 is the most likely where 1q gain and HY occured at the same time.
