## Supplementary Data 2 for "Temporal Genomic Dynamics Shape Clinical Trajectory in Multiple Myeloma"

### Timing the first multi-gain events in Multiple Myeloma

#### Introduction: Molecular and absolute time in Multiple Myeloma

Most tumors and virtually all normal tissues accumulate mutations at a consistent rate, typically specific to each tissue (Alexandrov et al., Nat Gen 2015; Gerstung et al., Nature 2020; Moore et al., Nature 2021). This accumulation rate, often referred to as a biological ‘clock,’ is minimally influenced by the tissue’s replication rate (Abascal et al. Nature 2021). The estimation of this clock relies on two distinct single base substitution (SBS) signatures: SBS1 and SBS5. Notably, in both aggressive and indolent lymphoma, as well as in normal B-cells, SBS1 and SBS5 exhibit a linear correlation with patients’ age (Gerstung et al., Nature 2020; Machado et al., Nature 2020), thereby confirming the presence of the biological clock, even in the context of germinal center exposure.

Earlier studies, albeit on limited sample sizes, indicated a similar clock correlation in multiple myeloma (Rustad et al., Nat Comm 2020). Building upon these preliminary findings, our study delved into 421 whole-genome sequencing (WGS) samples from multiple myeloma patients to validate and expand upon these early observations. We demonstrate that the mutational burden represented by SBS1 and SBS5 can effectively serve as a temporal marker, aligning with findings in other cancer types.”

I refined the language for clarity and coherence, ensured proper citation formatting, and provided a smoother transition between sentences. Let me know if you need further adjustments!

```
packages <- c('readr', 'tidyr', "splitstackshape", "plyr", "dplyr", "ggplot2", "ggpubr",
"reshape2", "magrittr", "lme4", "lmerTest", "knitr",
              "ggplot2","reshape2", "MASS", "RColorBrewer", "stringr", "deconstructSig
s", "BSgenome.Hsapiens.UCSC.hg19",
              "stringi", "tibble", "pander", "RColorBrewer", "merTools")

invisible(suppressWarnings(suppressMessages(lapply(packages, library, character.only = T
RUE))))
```

The uploaded file contains the contribution of each SBS signature for each sample. SBS signatures were estimated using mmsig (https://github.com/UM-Myeloma-Genomics/mmsig (https://github.com/UM-Myeloma-Genomics/mmsig))

```
# upload file with the mutational signature contribution for each patient
sig_age<- read.delim("timing_clock.txt")
head(sig_age)
```

```
##                 sample    SBS1    SBS2   SBS5   SBS8   SBS9 SBS13   SBS18
## 1 H130587_T02_02_WG01 0.07646 0.02476 0.4324 0.1237 0.3426     0 0.00000
## 2 H130608_T02_02_WG01 0.06264 0.02798 0.4687 0.1985 0.2422     0 0.00000
## 3 H130672_T03_01_WG01 0.03015 0.04871 0.4161 0.2263 0.2787     0 0.00000
## 4 H135228_T01_02_WG01 0.03034 0.03549 0.3865 0.1558 0.3919     0 0.00000
## 5 H135244_T01_02_WG01 0.06439 0.00000 0.3607 0.1964 0.3785     0 0.00000
## 6 H135256_T02_01_WG01 0.02118 0.00000 0.4379 0.2784 0.1934     0 0.06917
##   SBS.MM1 SBS31 SBS35 mutations IGH_partner    Ig chromosome HRD_status
## 1       0     0     0      3121       CCND1   IGH         11         no
## 2       0     0     0      6837         MYC IgK|L          8        yes
## 3       0     0     0      7095       CCND1   IGH         11         no
## 4       0     0     0      5198        NSD2   IGH          4        yes
## 5       0     0     0      4186       CCND1   IGH         11         no
## 6       0     0     0     14471         MYC IgK|L          8        yes
##   all.info.present          group                notes x1q HRD_window
## 1              yes      IGH_canon no gain with IGH TRA  no       <NA>
## 2              yes IGKL_other_HRD  focal TI on diploid  no      early
## 3              yes      IGH_canon no gain with IGH TRA yes       <NA>
## 4              yes  IGH_canon_HRD                 <NA> yes      early
## 5              yes      IGH_canon no gain with IGH TRA  no       <NA>
## 6              yes IGKL_other_HRD                 <NA> yes      early
##   IGH_window x1q_window HRD_moltime IGH_moltime x1q_mol_time_1 x1q_mol_time_2
## 1       <NA>       <NA>          NA          NA             NA             NA
## 2       <NA>       <NA>      0.6621          NA             NA             NA
## 3       <NA>      early          NA          NA             NA             NA
## 4      early      early      0.6614        0.84           0.68             NA
## 5       <NA>       <NA>          NA          NA             NA             NA
## 6       <NA>       <NA>      0.4692          NA             NA             NA
##            TRA_time x1q_SCT HRD_SCT IGH_SCT HRD_chrom_earliest_gain
## 1              <NA>    <NA>    <NA>    <NA>                    <NA>
## 2              <NA>    <NA>    <NA>    <NA>                    chr9
## 3              <NA>    <NA>    <NA>    <NA>                    <NA>
## 4       before gain    <NA>    <NA>    <NA>                   chr15
## 5              <NA>    <NA>    <NA>    <NA>                    <NA>
## 6 unrelated to gain    <NA>    <NA>    <NA>                   chr19
##   HRD_early_chrom_moltime           genomic_cluster    ASCT disease_status
## 1                      NA             CCND1_Simple No_ASCT           NDMM
## 2                  0.5810                HRD_Gains No_ASCT           NDMM
## 3                      NA CCND1_Complex_Cytogenetic No_ASCT           NDMM
## 4                  0.5656                 NSD2_HRD  ASCT_1           NDMM
## 5                      NA             CCND1_Simple No_ASCT           NDMM
## 6                  0.3137   HRD_Complex_Cytogenetic No_ASCT           NDMM
##   age_at_diagnosis age_at_sample_collection cohort patient abs_clock  stage_col
## 1               63                       63   DKRd H130587      1588 firebrick2
## 2               68                       68   DKRd H130608      3633 firebrick2
## 3               67                       67   DKRd H130672      3166 firebrick2
## 4               56                       56   DKRd H135228      2167 firebrick2
## 5               62                       62   DKRd H135244      1780 firebrick2
## 6               50                       50   DKRd H135256      6643 firebrick2
##   apobec_hyper apo_rr_state
## 1  deepskyblue         NDMM
## 2  deepskyblue         NDMM
```

```
## 3  deepskyblue         NDMM
## 4  deepskyblue         NDMM
## 5  deepskyblue         NDMM
## 6  deepskyblue         NDMM
```

Upload purity and coverage data and combine with mutational signature data frame

```
coverage_purity<- read.delim("Pre_Post_gain_JCOclassification_purity_coverage_421WGS.tx
t")
sig_age[is.na(sig_age)]<-""
coverage_purity[is.na(coverage_purity)]<-""
sig_age_cov<- unique(merge(sig_age, coverage_purity, by="sample"))
sig_age_cov$age_at_sample_collection<- as.numeric(as.character(sig_age_cov$age_at_sample
_collection)) # age at sample collection
sig_age_cov$abs_clock<- as.numeric(as.character(sig_age_cov$abs_clock)) # number of SBS1
and SBS5 mutations
sig_plot<- sig_age_cov[order(sig_age_cov$apo_rr_state),]
sig_plot<- unique(sig_plot[!is.na(sig_plot$age_at_sample_collection),])
```

Similarly to Gerstung et al. we assessed if the the relationship between the mutational burden of SBS1 and SBS5 and the age at sample collection could be explained by a linear model with the intercept constrained to zero.

```
### Interception constrained to zero as Gerstung et al. Nature 2020

summary(lm(sig_plot$abs_clock~
             0+sig_plot$age_at_sample_collection))
```

```
##
## Call:
## lm(formula = sig_plot$abs_clock ~ 0 + sig_plot$age_at_sample_collection)
##
## Residuals:
##    Min     1Q Median     3Q    Max
##  -3141   -805   -243    610   4532
##
## Coefficients:
##                                   Estimate Std. Error t value
## sig_plot$age_at_sample_collection    50.52       1.04    48.5
##                                               Pr(>|t|)
## sig_plot$age_at_sample_collection <0.0000000000000002 ***
## ---
## Signif. codes:  0 '***' 0.001 '**' 0.01 '*' 0.05 '.' 0.1 ' ' 1
##
## Residual standard error: 1270 on 416 degrees of freedom
## Multiple R-squared:  0.85,   Adjusted R-squared:  0.849
## F-statistic: 2.35e+03 on 1 and 416 DF,  p-value: <0.0000000000000002
```

Then, the regression model was adjusted for hyper-APOBEC activity, the presence of melphalan mutational signatures, and disease stage (relapse vs. newly diagnosed).

```
summary(lm((sig_plot$abs_clock~sig_plot$age_at_sample_collection+sig_plot$apo_rr_stat
e)))
```

```
##
## Call:
## lm(formula = (sig_plot$abs_clock ~ sig_plot$age_at_sample_collection +
##     sig_plot$apo_rr_state))
##
## Residuals:
##    Min     1Q Median     3Q    Max
##  -4228   -695    -63    499   4240
##
## Coefficients:
##                                            Estimate Std. Error t value
## (Intercept)                                 2957.51     405.52    7.29
## sig_plot$age_at_sample_collection             23.41       6.32    3.71
## sig_plot$apo_rr_stateNDMM                  -1725.17     151.23  -11.41
## sig_plot$apo_rr_stateRRMM with Melphalan     -911.56     200.90   -4.54
## sig_plot$apo_rr_stateRRMM without Melphalan -1539.34     271.03   -5.68
##                                                         Pr(>|t|)
## (Intercept)                                      0.0000000000016 ***
## sig_plot$age_at_sample_collection                        0.00024 ***
## sig_plot$apo_rr_stateNDMM                   < 0.0000000000000002 ***
## sig_plot$apo_rr_stateRRMM with Melphalan         0.0000074815468 ***
## sig_plot$apo_rr_stateRRMM without Melphalan      0.0000000255362 ***
## ---
## Signif. codes:  0 '***' 0.001 '**' 0.01 '*' 0.05 '.' 0.1 ' ' 1
##
## Residual standard error: 1100 on 412 degrees of freedom
## Multiple R-squared:  0.29,   Adjusted R-squared:  0.284
## F-statistic: 42.2 on 4 and 412 DF,  p-value: <0.0000000000000002
```

Visualize the correlation between SBS1 and SBS5 and age in multiple myeloma

```
### ALL SBS5 and SBS1
par(xpd=F, mar=c(5,5,2,5))
plot(x=sig_plot$age_at_sample_collection,pch=21,bg=sig_plot$apobec_hyper,
     y=sig_plot$abs_clock, xlim=c(0, 100), ylim=c(0,10000), las=2,
     yaxt="n", ylab="", xlab="", xaxt="n", bty="n")
axis(side = 2, at = seq(0,10000, by=2000),labels = seq(0,10000, by=2000), las=2, cex.axi
s=1.5, lwd=1.5)
axis(side = 1, at = seq(0,100, by=20),labels = seq(0,100, by=20), las=1, cex.axis=1.5, l
wd=1.5)
abline(lm(sig_plot$abs_clock~sig_plot$age_at_sample_collection+sig_plot$apobec_hyper))
```

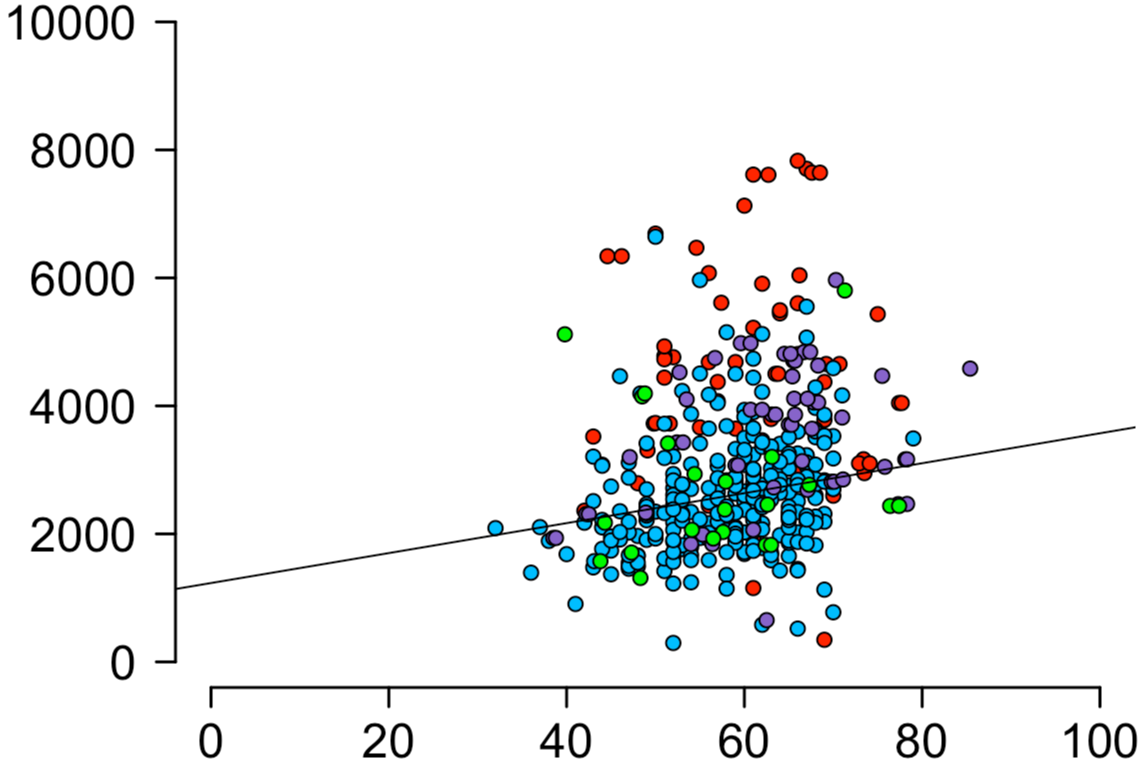


We further investigate the correlation between SBS1/SBS5 and age while correcting for potential confounders such as coverage and purity Additionally, the presence of hyperAPOBEC was tested, as we previously demonstrated that, similar to other hypermutated tumors (e.g., Melanoma), the clock-like signal becomes unreliable due to the interbreeding of the mutations process causing hypermutation.

```
### coverage and purity don;t affect the correlation

sig_plot$Coverage.TUMOR<- as.numeric(as.character(sig_plot$Coverage.TUMOR))
sig_plot$Purity<- as.numeric(as.character(sig_plot$Purity))

summary(lm(sig_plot$abs_clock~
             sig_plot$age_at_sample_collection + sig_plot$apo_rr_state+
               sig_plot$Coverage.TUMOR/sig_plot$Purity))
```

```
##
## Call:
## lm(formula = sig_plot$abs_clock ~ sig_plot$age_at_sample_collection +
##     sig_plot$apo_rr_state + sig_plot$Coverage.TUMOR/sig_plot$Purity)
##
## Residuals:
##    Min     1Q Median     3Q    Max
##  -3963   -596    -76    471   4366
##
## Coefficients:
##                                           Estimate Std. Error t value
## (Intercept)                                2051.75     572.99    3.58
## sig_plot$age_at_sample_collection            26.08       7.45    3.50
## sig_plot$apo_rr_stateNDMM                 -1436.94     171.11   -8.40
## sig_plot$Coverage.TUMOR                       2.89       5.63    0.51
## sig_plot$Coverage.TUMOR:sig_plot$Purity       2.89       3.93    0.73
##                                                     Pr(>|t|)
## (Intercept)                                          0.00040 ***
## sig_plot$age_at_sample_collection                    0.00053 ***
## sig_plot$apo_rr_stateNDMM                 0.0000000000000017 ***
## sig_plot$Coverage.TUMOR                              0.60844
## sig_plot$Coverage.TUMOR:sig_plot$Purity              0.46303
## ---
## Signif. codes:  0 '***' 0.001 '**' 0.01 '*' 0.05 '.' 0.1 ' ' 1
##
## Residual standard error: 1030 on 309 degrees of freedom
##   (103 observations deleted due to missingness)
## Multiple R-squared:  0.227,  Adjusted R-squared:  0.217
## F-statistic: 22.7 on 4 and 309 DF,  p-value: <0.0000000000000002
```

Here we estimate the residual for each patient. Patients with an excess of residual. To be consistent with recent litterature we decide to use the model with the intercept constrain to 0.

```
model_res<- (lm(sig_plot$abs_clock~ 0+sig_plot$age_at_sample_collection))
residuals_vec <- residuals(model_res)

# Calculate absolute values of residuals
abs_residuals <- abs(residuals_vec)
sig_plot$residual<- abs_residuals
sig_plot<- sig_plot[order(sig_plot$residual),]
plot((sig_plot$residual), col=sig_plot$apobec_hyper,pch=16, ylim=c(0,10000), ylab="Resid
ual")
```

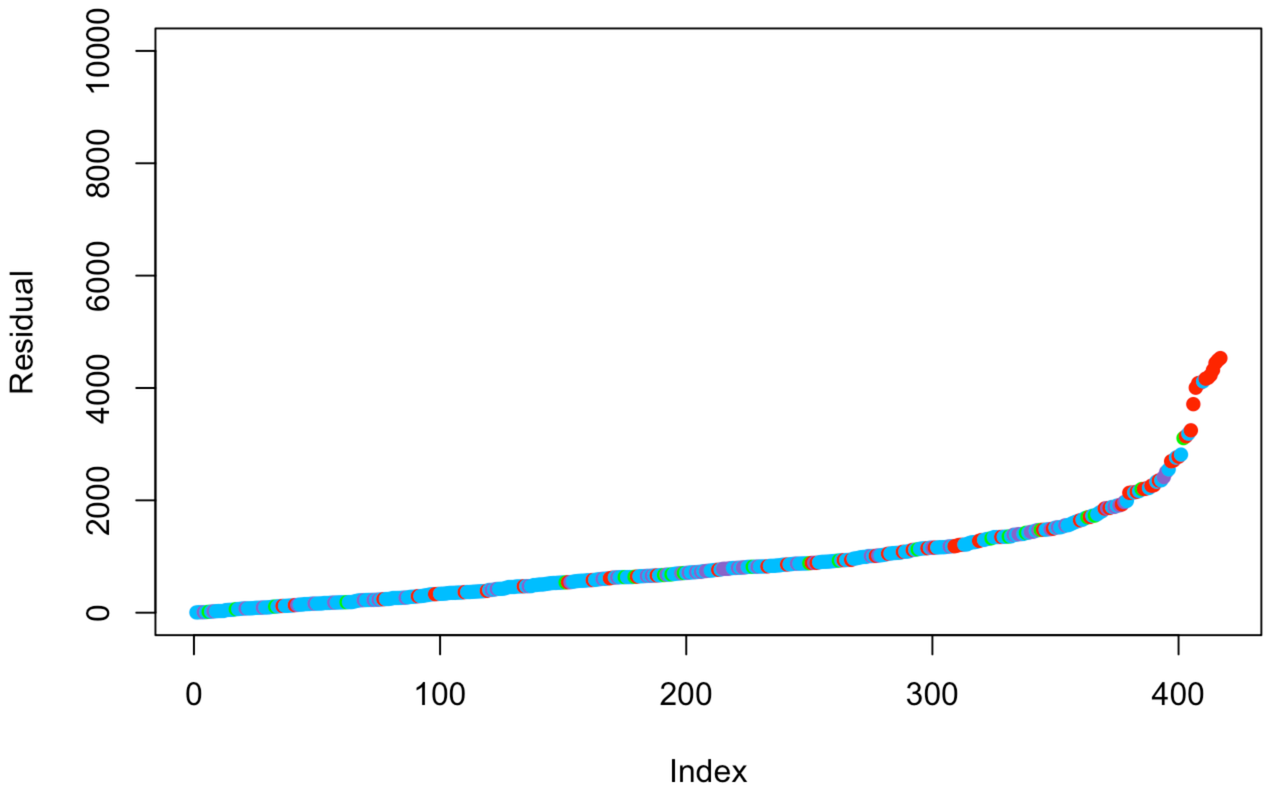


The number of cases that do not fit the linear model will be excluded from the absolute timing estimation analysis.

```
cutff_model<-1900
length(sig_plot$apobec_hyper[sig_plot$residual>cutff_model])
```

```
## [1] 43
```

```
sig_plot$residual_code<- 0
sig_plot$residual_code[sig_plot$residual>cutff_model]<- 1

par(xpd=F, mar=c(5,5,2,5))
# [sig_age2$abs_clock<8000,]
plot(sig_plot$age_at_sample_collection[sig_plot$residual>cutff_model],sig_plot$abs_clock
[sig_plot$residual>cutff_model],
     pch=16, col=alpha(sig_plot$apobec_hyper[sig_plot$residual>cutff_model], 0.6),
     xlim=c(0, 100), ylim=c(0,10000), las=2,
     yaxt="n", ylab="", xlab="", xaxt="n", bty="n")
par(new=T)
plot(sig_plot$age_at_sample_collection[sig_plot$residual>cutff_model],sig_plot$abs_clock
[sig_plot$residual>cutff_model],
     pch=21, bg = NA, col="grey",
     xlim=c(0, 100), ylim=c(0,10000), las=2,
     yaxt="n", ylab="", xlab="", xaxt="n", bty="n")

par(new=T)
plot(sig_plot$age_at_sample_collection[sig_plot$residual<cutff_model],pch=21,bg=sig_plot
$apobec_hyper,
     sig_plot$abs_clock[sig_plot$residual<cutff_model], xlim=c(0, 100), ylim=c(0,10000),
las=2,
     ylab="", xlab="",  bty="n")
abline(lm(sig_plot$abs_clock[sig_plot$residual<cutff_model]~0+sig_plot$age_at_sample_col
lection
          [sig_plot$residual<cutff_model]))
```

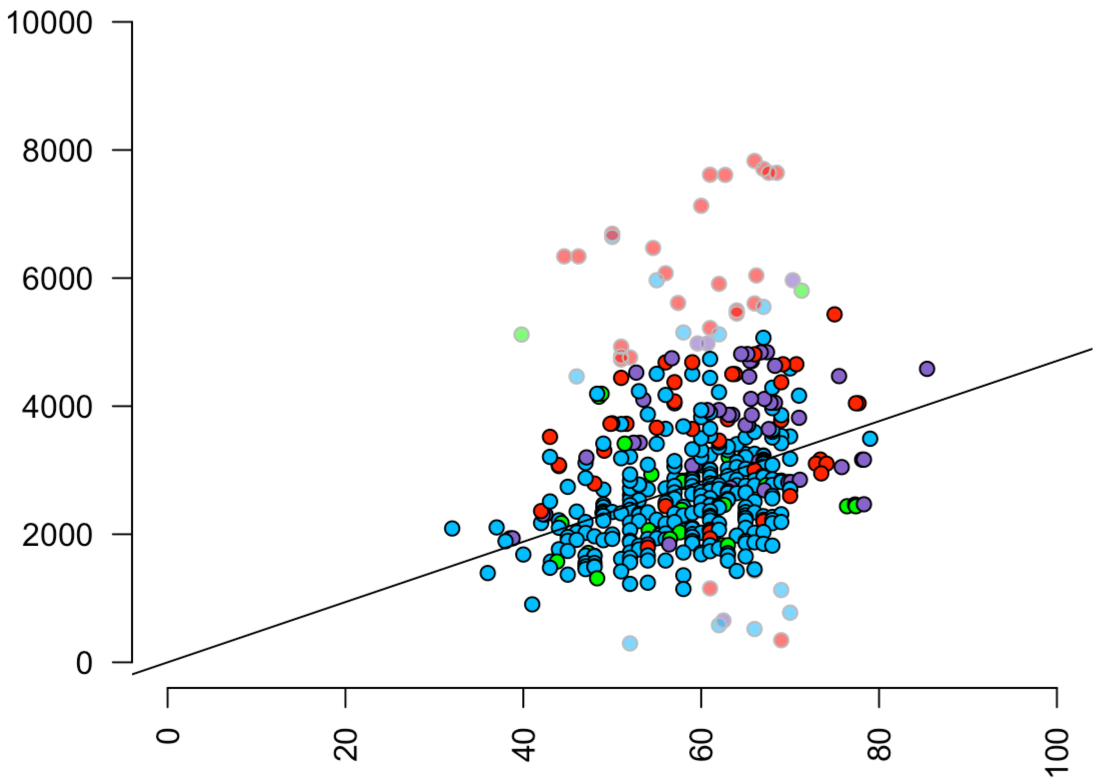


Comparing SBS1 and SBS5 mutation rate between multiple myeloma and normal B-cell. Normal B-cell data were taken from Machado et al. Nature 2022. Because Machado et al. used single cell expansion, to make the comparison more reliable we only considered clonal mutations in multiple myeloma WGS (i.e. the most recent common ancestor).

```
pcawgs<- read.delim("PCAWG_sigProfiler_SBS_signatures_in_samples.csv", sep=",")
pcawgs_lymph<-pcawgs[pcawgs$Cancer.Types %in% c("Lymph-BNHL"    ,      "Lymph-CLL" ),]

age_cpag<-read.delim("PCAWG7_age_information.txt")
colnames(age_cpag)<-c("Sample.Names","age")
pcag<- merge(age_cpag, pcawgs_lymph, by="Sample.Names")
pcag$clock<-pcag$SBS1+pcag$SBS5
pcag<-pcag[pcag$clock<8000,] ## remove hypermutated
pcag<-pcag[-grep("CLL", pcag$Cancer.Types),] ## remove CLL

sig_plot<- sig_plot[sig_plot$abs_clock<8000,] ## remove hypermutated

par(xpd=F, mar=c(5,5,2,5))
plot(sig_plot$age_at_sample_collection,pch=21,bg="dodgerblue",
     sig_plot$abs_clock, xlim=c(0, 100), ylim=c(0,10000), las=2,
     yaxt="n", ylab="", xlab="", xaxt="n", bty="n")

summary(lm((sig_plot$abs_clock~sig_plot$age_at_sample_collection+sig_plot$stage_col+sig_
plot$apobec_hyper)))
```

```
##
## Call:
## lm(formula = (sig_plot$abs_clock ~ sig_plot$age_at_sample_collection +
##     sig_plot$stage_col + sig_plot$apobec_hyper))
##
## Residuals:
##    Min     1Q Median     3Q    Max
##  -4035   -692    -92    529   4229
##
## Coefficients:
##                                    Estimate Std. Error t value
## (Intercept)                         1912.76     501.30    3.82
## sig_plot$age_at_sample_collection     21.96       6.33    3.47
## sig_plot$stage_colfirebrick2        -595.79     294.69   -2.02
## sig_plot$apobec_hypergreen          -411.56     382.14   -1.08
## sig_plot$apobec_hypermediumpurple3   224.90     334.38    0.67
## sig_plot$apobec_hyperred            1548.18     174.25    8.88
##                                                Pr(>|t|)
## (Intercept)                                     0.00016 ***
## sig_plot$age_at_sample_collection               0.00058 ***
## sig_plot$stage_colfirebrick2                    0.04385 *
## sig_plot$apobec_hypergreen                      0.28211
## sig_plot$apobec_hypermediumpurple3              0.50160
## sig_plot$apobec_hyperred           < 0.0000000000000002 ***
## ---
## Signif. codes:  0 '***' 0.001 '**' 0.01 '*' 0.05 '.' 0.1 ' ' 1
##
## Residual standard error: 1090 on 411 degrees of freedom
## Multiple R-squared:  0.297,  Adjusted R-squared:  0.289
## F-statistic: 34.8 on 5 and 411 DF,  p-value: <0.0000000000000002
```

```
abline(lm(sig_plot$abs_clock~sig_plot$age_at_sample_collection+sig_plot$apobec_hyper), c
ol="dodgerblue4")
```

```
## Warning in abline(lm(sig_plot$abs_clock ~ sig_plot$age_at_sample_collection + :
## only using the first two of 5 regression coefficients
```

```
axis(side = 2, at = seq(0,10000, by=2000),labels = seq(0,10000, by=2000), las=2, cex.axi
s=1.5, lwd=1.5)
axis(side = 1, at = seq(0,100, by=20),labels = seq(0,100, by=20), las=1, cex.axis=1.5, l
wd=1.5)
par(new=T)
plot(x=pcag$age,pch=21,bg="olivedrab3",
     y=pcag$clock, xlim=c(0, 100), ylim=c(0,10000), las=2,
     yaxt="n", ylab="", xlab="", xaxt="n", bty="n", col="black")
abline(lm(pcag$clock~pcag$age), col="forestgreen")
```

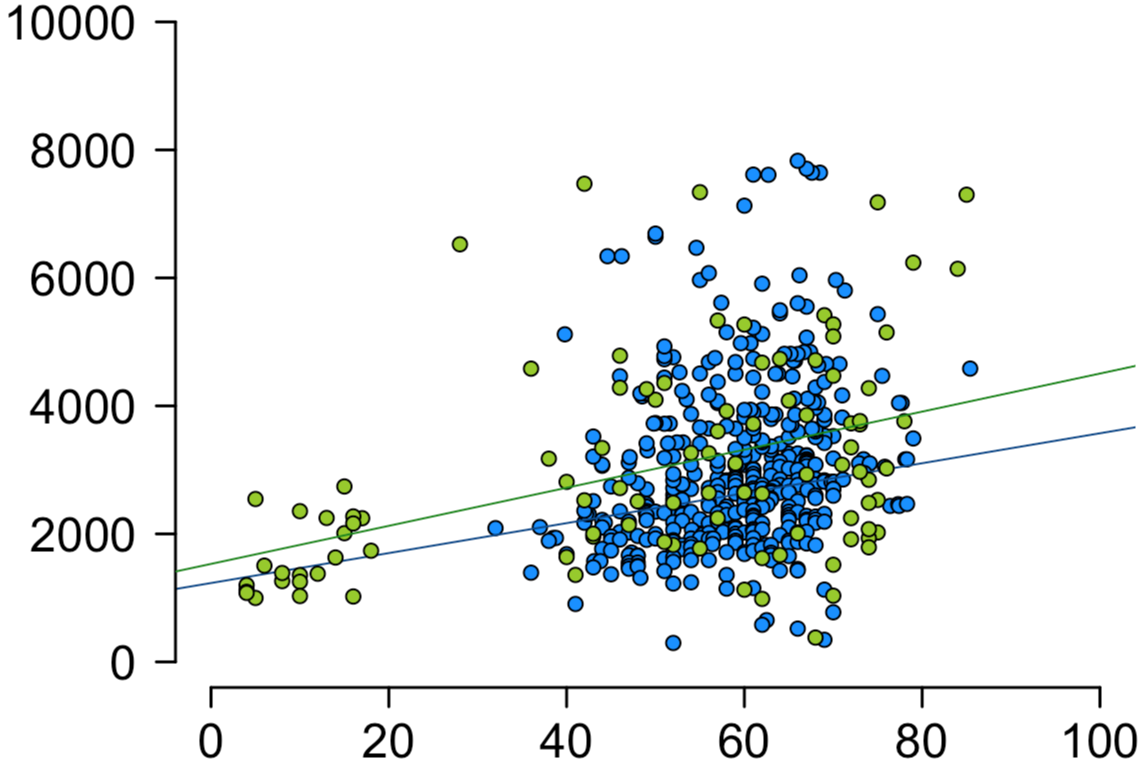


```
summary(lm(pcag$clock~pcag$age))
```

```
##
## Call:
## lm(formula = pcag$clock ~ pcag$age)
##
## Residuals:
##    Min     1Q Median     3Q    Max
##  -3175   -820   -287    687   4690
##
## Coefficients:
##             Estimate Std. Error t value  Pr(>|t|)
## (Intercept)  1531.27     334.54    4.58 0.0000138 ***
## pcag$age       29.77       6.08    4.89 0.0000039 ***
## ---
## Signif. codes:  0 '***' 0.001 '**' 0.01 '*' 0.05 '.' 0.1 ' ' 1
##
## Residual standard error: 1470 on 98 degrees of freedom
## Multiple R-squared:  0.196,  Adjusted R-squared:  0.188
## F-statistic:   24 on 1 and 98 DF,  p-value: 0.00000388
```

```
summary(lm(pcag$clock~0+pcag$age)) # constrain intercept like Gerstung et al. Nature 202
0
```

```
##
## Call:
## lm(formula = pcag$clock ~ 0 + pcag$age)
##
## Residuals:
##    Min     1Q Median     3Q    Max
##  -3344   -779    481   1259   5171
##
## Coefficients:
##          Estimate Std. Error t value            Pr(>|t|)
## pcag$age    54.77       2.93    18.7 <0.0000000000000002 ***
## ---
## Signif. codes:  0 '***' 0.001 '**' 0.01 '*' 0.05 '.' 0.1 ' ' 1
##
## Residual standard error: 1610 on 99 degrees of freedom
## Multiple R-squared:  0.779,  Adjusted R-squared:  0.777
## F-statistic:  349 on 1 and 99 DF,  p-value: <0.0000000000000002
```

```
machado<- read.delim("machado_mmsig.txt")
machado$id <- as.factor(machado$id)
collapsed_df <- aggregate(abs_clock ~ id, data = machado, FUN = median, na.rm = TRUE)
collapsed_df <- merge(collapsed_df, machado[!duplicated(machado$id), c("id", "Age", "Cel
lType", "Cell.type2", "Tissue")], by = "id")
collapsed_df2<- collapsed_df[collapsed_df$Cell.type2=="Memory B",]


clonal_sig2<- read.delim("CLONAL_SBS.txt",sep="\t")


par(xpd=F, mar=c(5,5,2,5))
plot(clonal_sig2$age_at_sample_collection,pch=21,bg=clonal_sig2$apobec_hyper,
     clonal_sig2$abs_clock, xlim=c(0, 100), ylim=c(0,6000), las=2,
     yaxt="n", ylab="", xlab="", xaxt="n", bty="n")
axis(side = 2, at = seq(0,6000, by=2000),labels = seq(0,6000, by=2000), las=2, cex.axis=
1.5, lwd=1.5)
axis(side = 1, at = seq(0,100, by=20),labels = seq(0,100, by=20), las=1, cex.axis=1.5, l
wd=1.5)
par(new=T)
machado2<-machado[machado$CellType == "B Memory",]
plot(machado$Age, machado$abs_clock, pch=21,bg="gold3",
     xlim=c(0, 100), ylim=c(0,6000), las=2,
     yaxt="n", ylab="", xlab="", xaxt="n", bty="n")
abline(lm(machado$abs_clock~0+machado$Age),col = "black")
abline(lm(clonal_sig2$abs_clock~0+clonal_sig2$age_at_sample_collection),col = "red")
summary(lm(machado2$abs_clock~0+machado2$Age))
```

```
##
## Call:
## lm(formula = machado2$abs_clock ~ 0 + machado2$Age)
##
## Residuals:
##    Min     1Q Median     3Q    Max
## -830.7 -153.2   75.2  320.9 3005.9
##
## Coefficients:
##              Estimate Std. Error t value          Pr(>|t|)
## machado2$Age    17.29       1.57      11 0.000000000000043 ***
## ---
## Signif. codes:  0 '***' 0.001 '**' 0.01 '*' 0.05 '.' 0.1 ' ' 1
##
## Residual standard error: 626 on 43 degrees of freedom
## Multiple R-squared:  0.738,  Adjusted R-squared:  0.732
## F-statistic:  121 on 1 and 43 DF,  p-value: 0.0000000000000426
```

```
summary(lm(clonal_sig2$abs_clock~clonal_sig2$age_at_sample_collection+clonal_sig2$apobec
_hyper))
```

```
##
## Call:
## lm(formula = clonal_sig2$abs_clock ~ clonal_sig2$age_at_sample_collection +
##     clonal_sig2$apobec_hyper)
##
## Residuals:
##     Min      1Q  Median      3Q     Max
## -1750.1  -399.7   -43.6   342.3  2744.8
##
## Coefficients:
##                                       Estimate Std. Error t value  Pr(>|t|)
## (Intercept)                             530.91     234.83    2.26     0.024 *
## clonal_sig2$age_at_sample_collection     17.47       3.99    4.38 0.0000157 ***
## clonal_sig2$apobec_hypergreen           230.73     139.93    1.65     0.100
## clonal_sig2$apobec_hypermediumpurple3   428.20      95.25    4.50 0.0000095 ***
## ---
## Signif. codes:  0 '***' 0.001 '**' 0.01 '*' 0.05 '.' 0.1 ' ' 1
##
## Residual standard error: 631 on 347 degrees of freedom
##   (1 observation deleted due to missingness)
## Multiple R-squared:  0.127,  Adjusted R-squared:  0.119
## F-statistic: 16.8 on 3 and 347 DF,  p-value: 0.000000000346
```

```
legend("toprigh", legend = c("Normal B-cells", "MM"), pch = "-", col=c("black","red"))
```

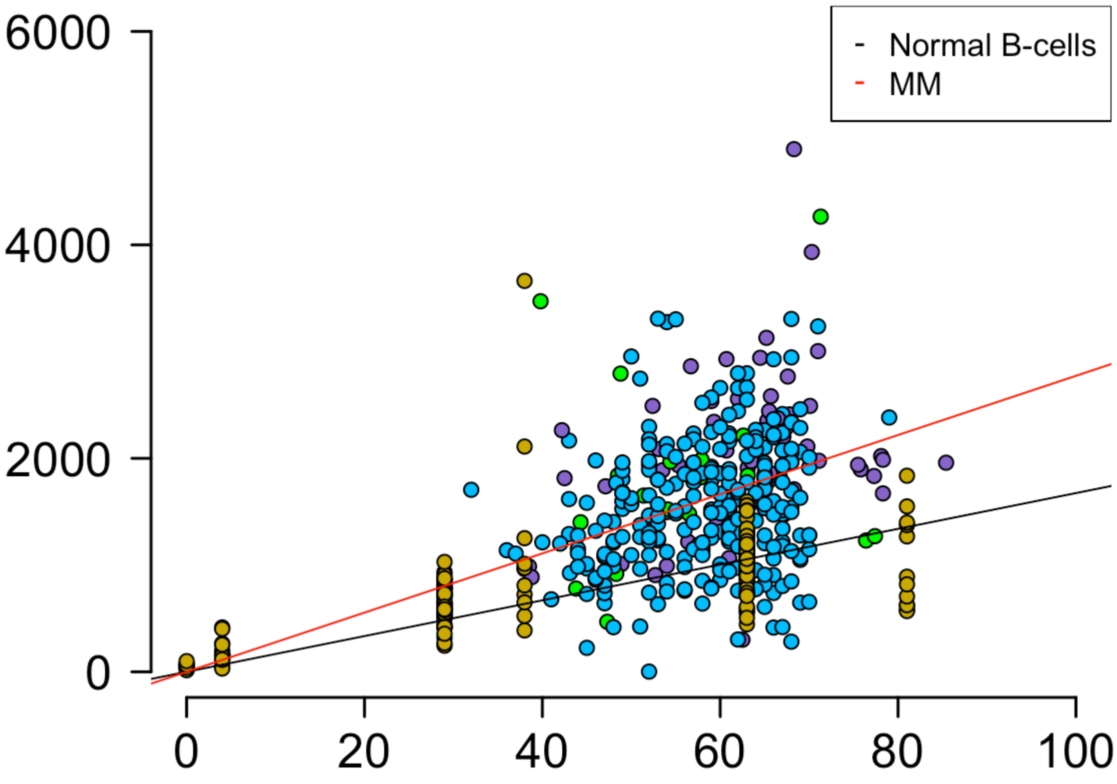


Compare the linear model from Machado et al. Nature 2020 B-cell SBS-SBS5 mutation rate and the one observed in multiple myeloma

```
model1 <- lm(machado2$abs_clock~0+machado2$Age)
model2 <- lm(clonal_sig2$abs_clock ~0+clonal_sig2$age_at_sample_collection+clonal_sig2$a
pobec_hyper)
coef_model1 <- coef(summary(model1))["machado2$Age", "Estimate"]
coef_model2 <- coef(summary(model2))["clonal_sig2$age_at_sample_collection", "Estimate"]
se_model1 <- coef(summary(model1))["machado2$Age", "Std. Error"]
se_model2 <- coef(summary(model2))["clonal_sig2$age_at_sample_collection", "Std. Error"]

# Calculate the t-value
t_value <- (coef_model1 - coef_model2) / sqrt(se_model1^2 + se_model2^2)

# Calculate the degrees of freedom
df <- sum(model1$df.residual, model2$df.residual)

# Perform a t-test
p_value <- 2 * pt(abs(t_value), df = df, lower.tail = FALSE)
print(p_value)
```

```
## [1] 0.9661
```
