## Supplementary figures and images for "Temporal Genomic Dynamics Shape Clinical Trajectory in Multiple Myeloma"

### Supplementary Data 3

MM024

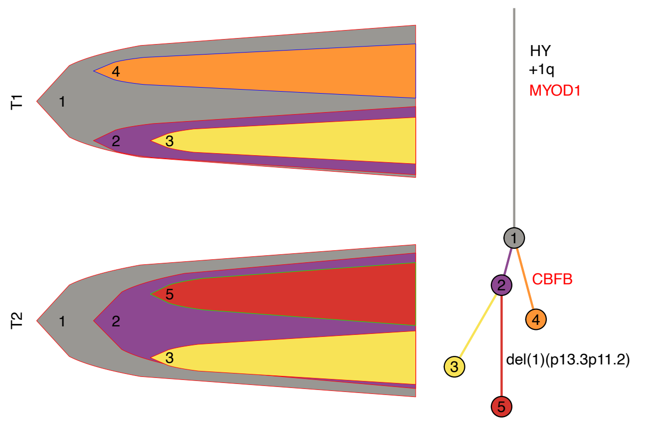


MM026

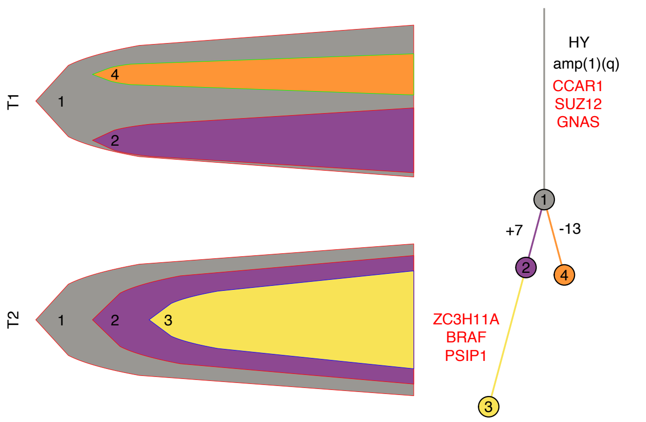


MM027

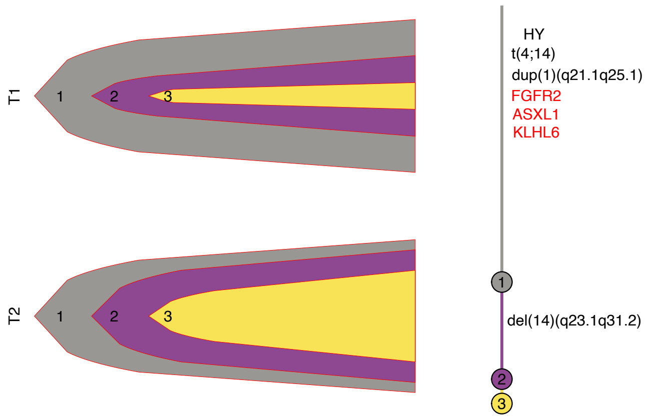


MM031

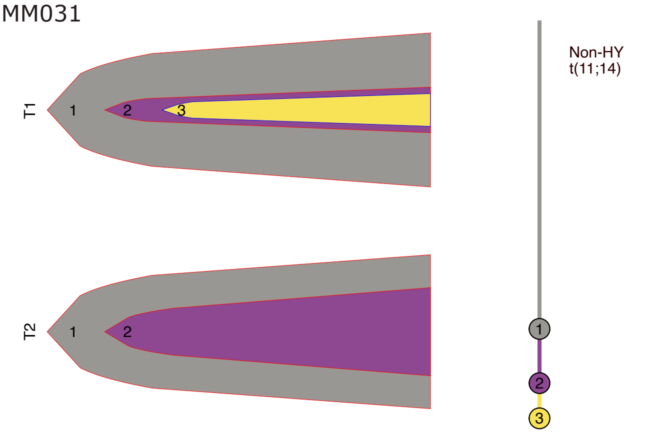


MM036

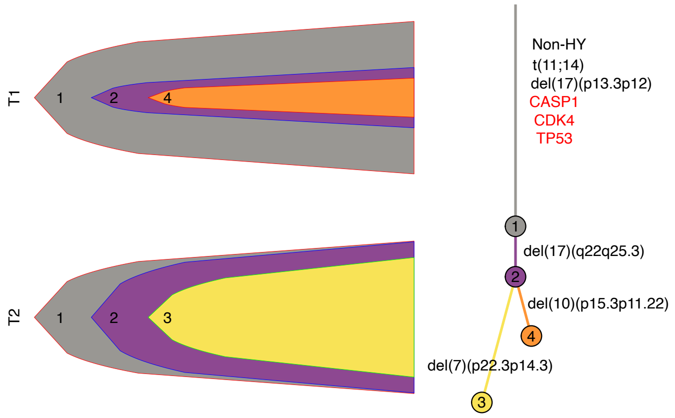


MM045

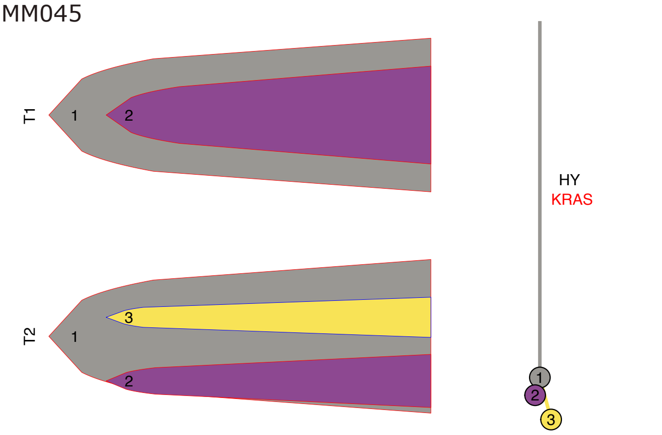


MM078

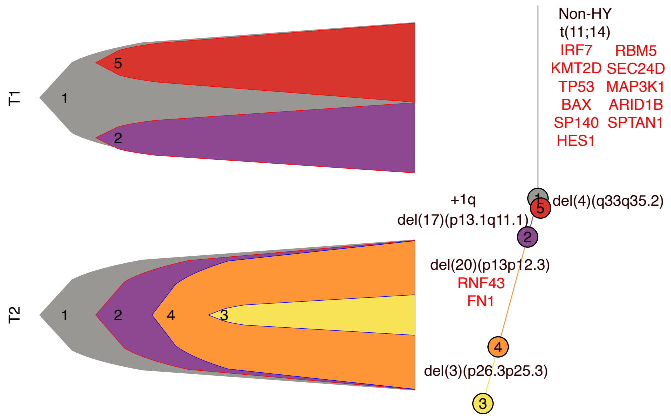


MM133

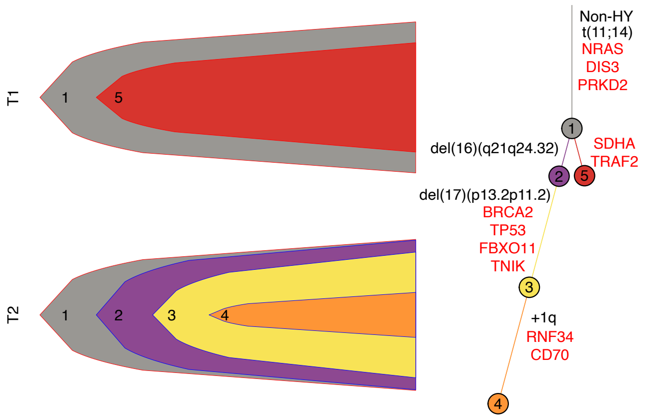

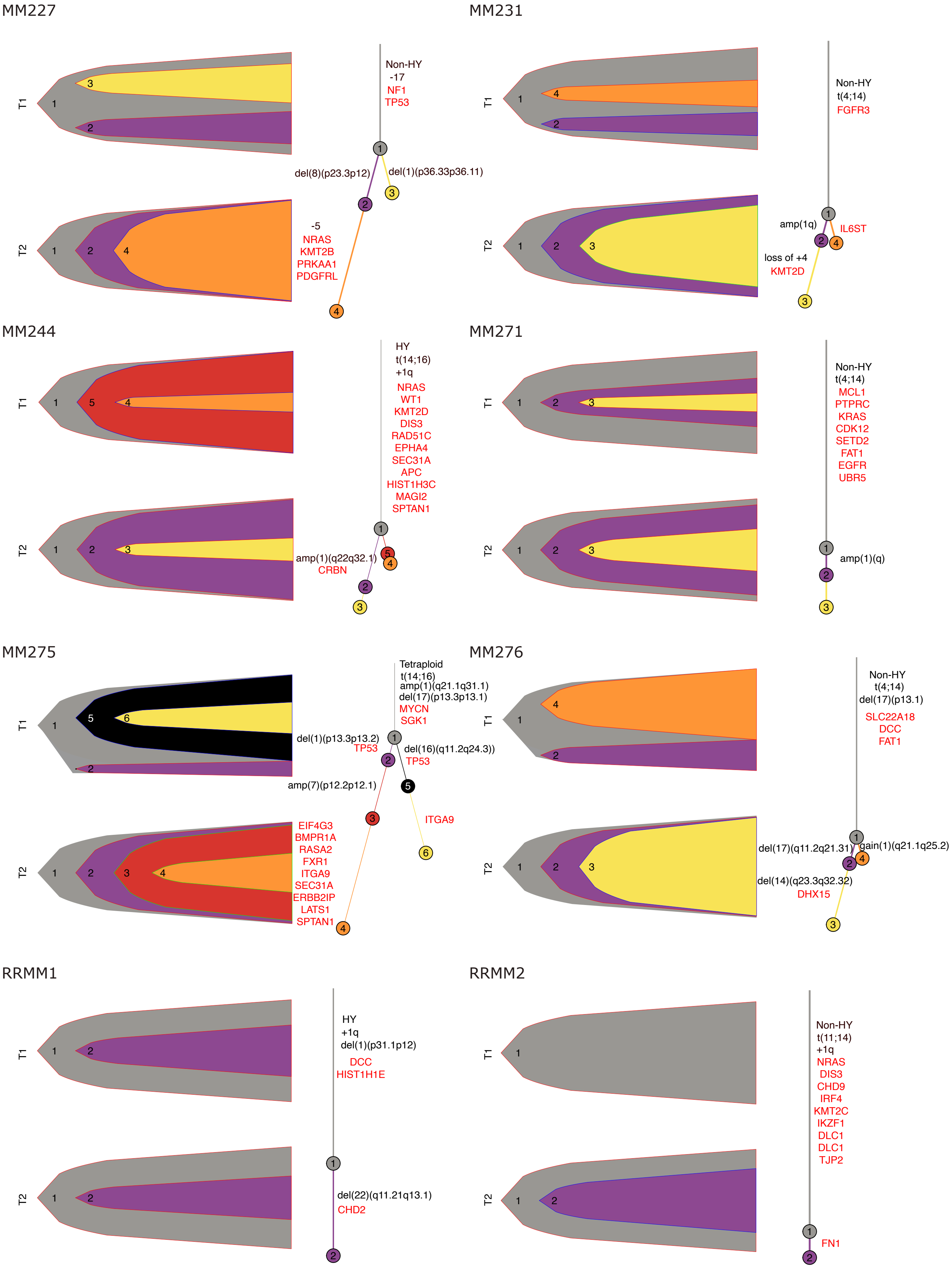

RRMM4

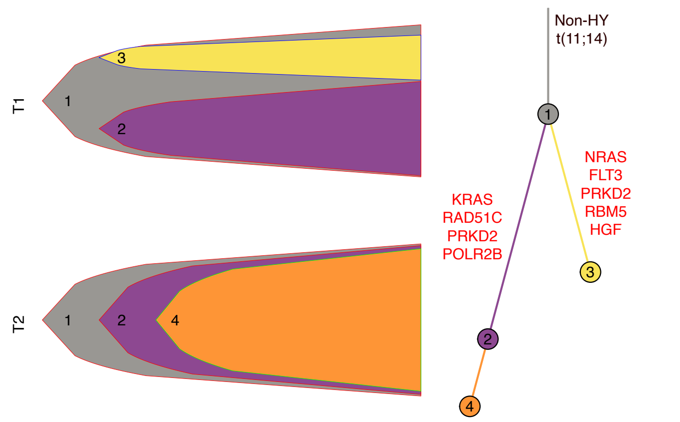


RRMM5

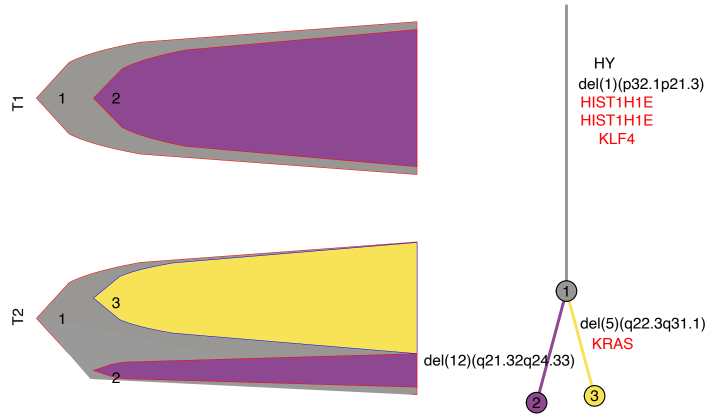


RRMM7

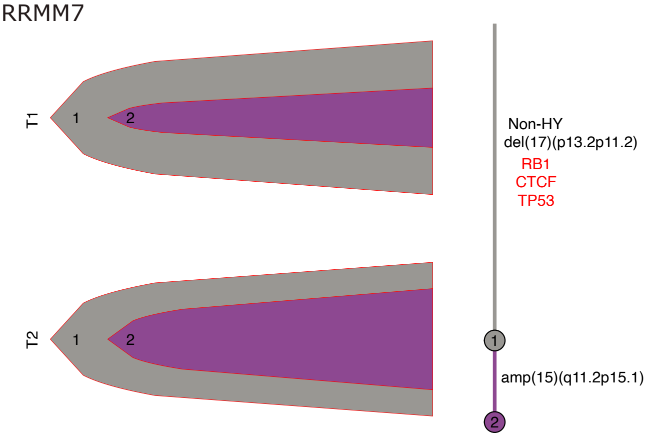


RRMM9

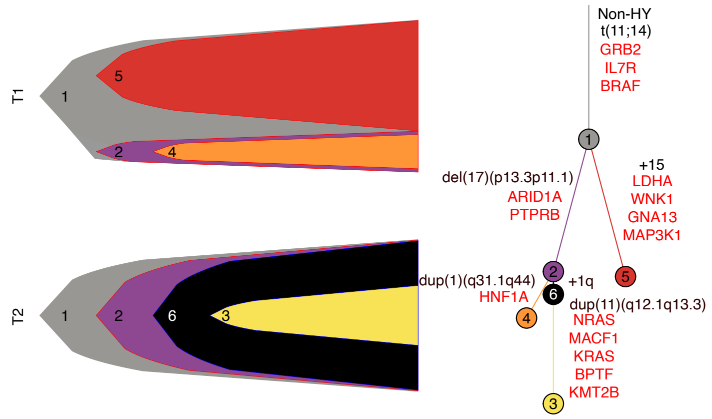


RRMM10

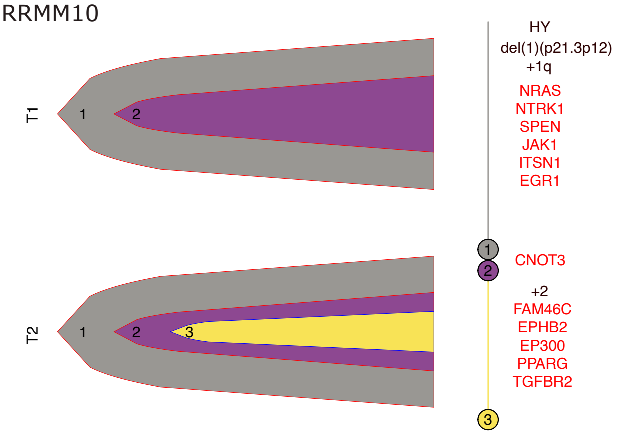


RRMM13

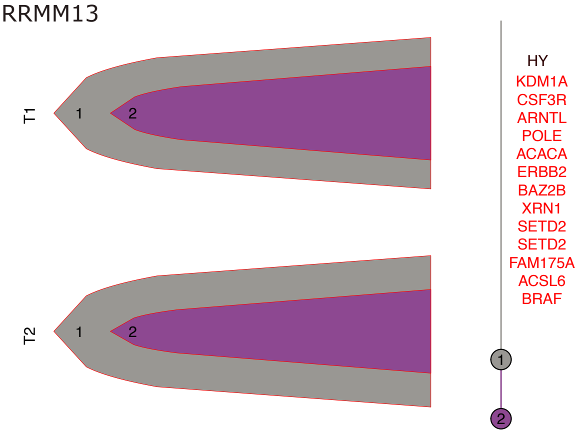


RRMM15

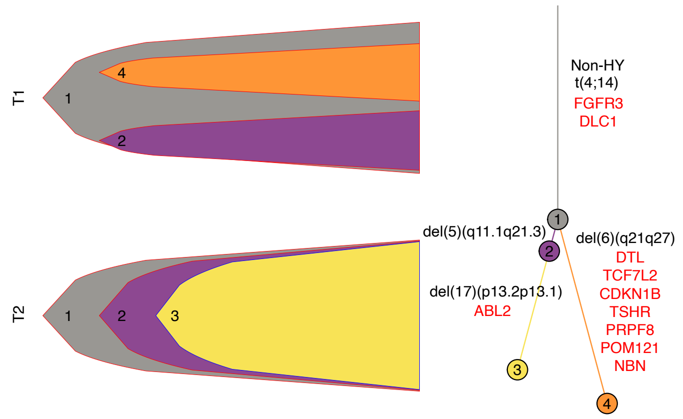


RRMM16

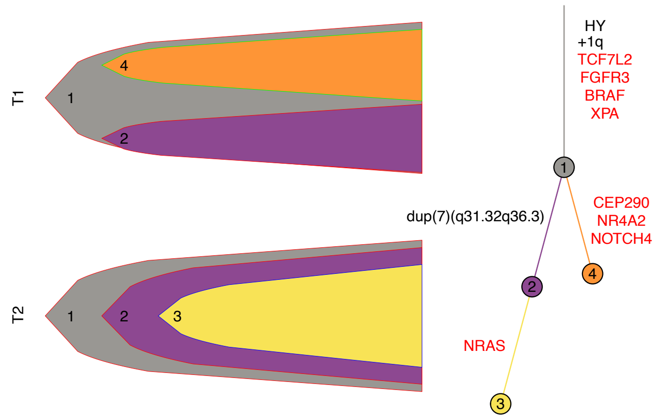

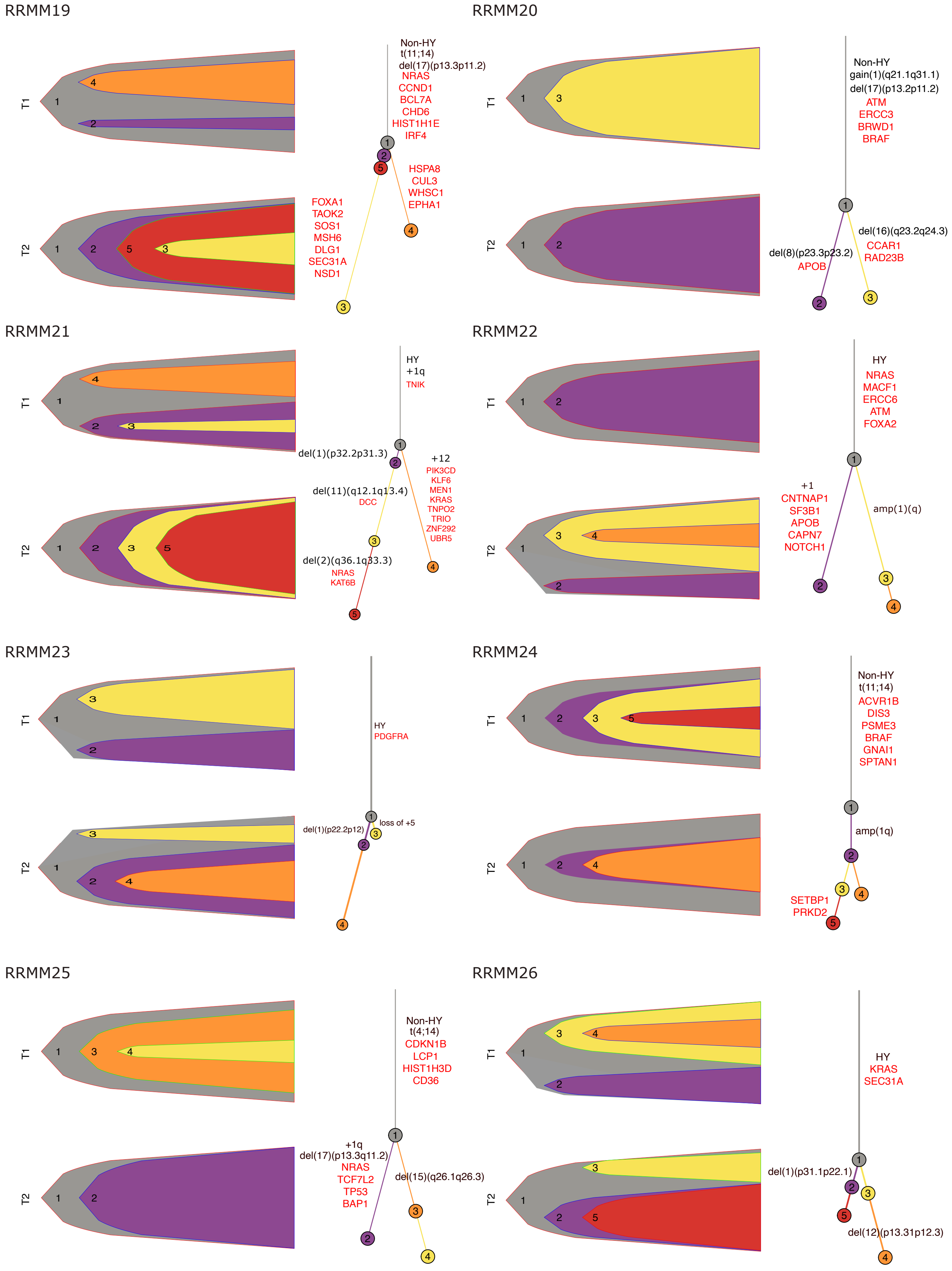

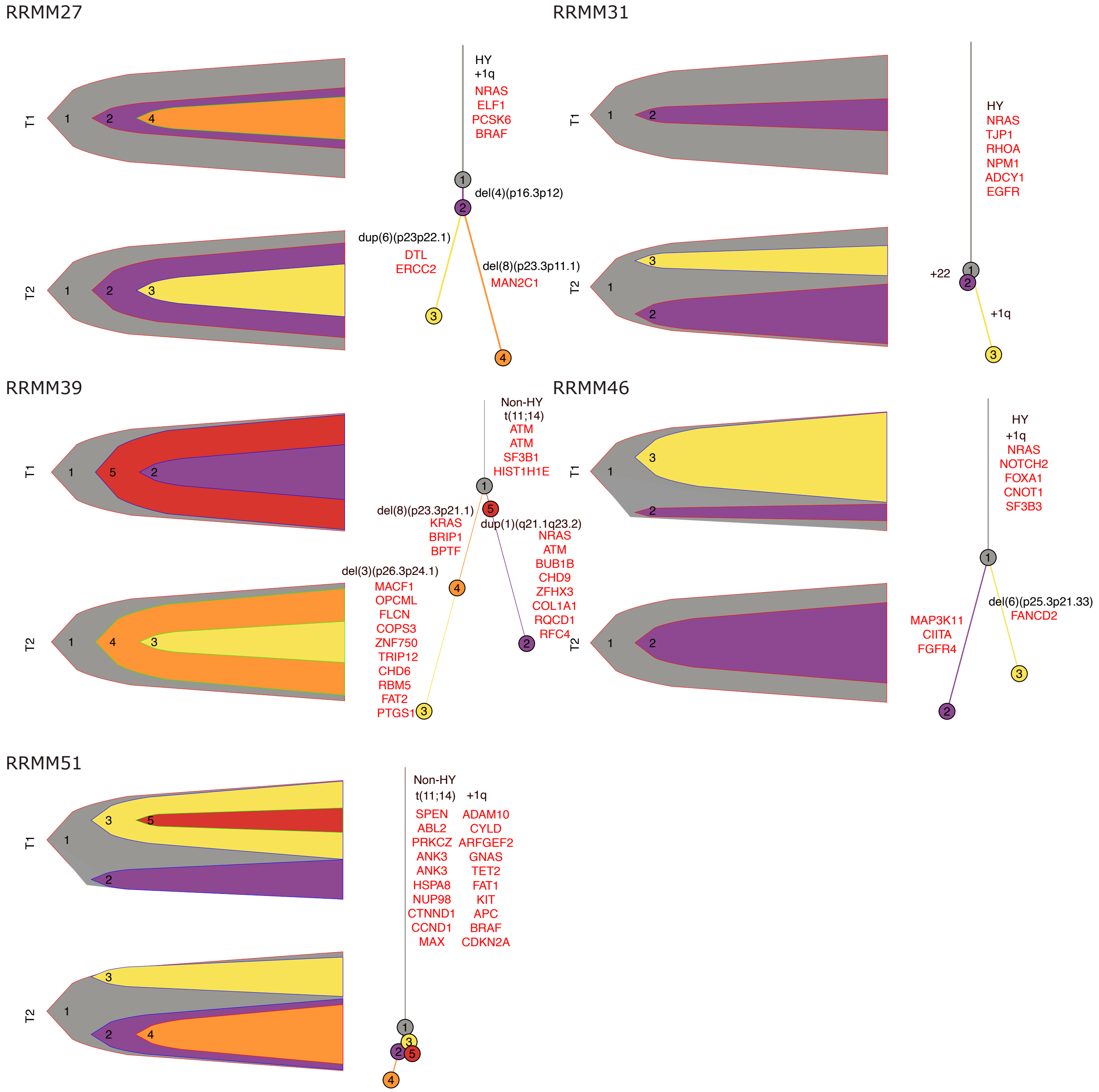
