## Supplementary Data 4 for "Temporal Genomic Dynamics Shape Clinical Trajectory in Multiple Myeloma"

### Supplementary Data 4 – Timing multiple myeloma evolution

Among patients with *MAF*/*MAFB* translocations and HY, we identified only one case (MM065) where 16q gain was linked to t(14;16)(*IGH*;*MAF*) (**Supplemental Data Fig. 1A**). Intriguingly, within this patient, we observed two distinct and significantly separated time frames: an early window associated with 16q gain, acquired after the t(14;16)(*IGH*;*MAF*) reciprocal translocation and a later one where the second 16q gain was linked to a multi-gain event with whole genome doubling (WGD), resulting in a tetraploid karyotype. Notably, the large temporal gap between these two events was driven by the effect of high APOBEC mutational activity in increasing the post-gain mutation burden (hyper-APOBEC; **Fig. 3**). Interestingly, APOBEC mutagenesis was also detectable in the 16q pre-gain mutations (**Fig. Supplemental Data Fig. 1B**). Overall, this distinct profile can be explained by a three-step model: 1) a t(14;16)(*IGH*;*MAF*) reciprocal translocation created a derivative chromosome, promoting APOBEC mutagenesis; 2) the derivative chromosome is duplicated shortly after the MAF translocation and after acquiring a small burden of APOBEC-mediated mutations; 3) as a consequence of a WGD event and an additional 16q gain (**Supplemental Data Fig. 1C**). Despite this observation was limited to one single patient, these data represent additional evidence supporting the model where patients with *MAF*/*MAFB* translocations experience high APOBEC mutagenic activity early in time (**Figure 3B-C**) [1,2]. 1q gain is present in ~70% of MM patients with a *MAF*/*MAFB* translocations [t(14;16) or t(14;20)][3-6]. We therefore sought to investigate whether 1q gain is an early or late event in these patients. Because *MAF*/*MAFB* induces the hyper-APOBEC state, we assumed that acquisition of hyper-APOBEC before the 1q gain would imply that the *MAF*/*MAFB* translocation precedes the 1q gain. We identified 8 patients in our study that had a *MAF*/*MAFB* translocation and a 1q gain with molecular time data available for analysis. In all 8 patients, hyper-APOBEC was detected among the duplicated mutations on 1q, suggesting the *MAF*/*MAFB* translocation always preceded the 1q gain.

*NSD2* translocations [i.e., t(4;14)] predominantly result in deletions on chromosome 4p rather than gains. To estimate the timing of t(4;14)(*NSD2;IGH*), we employed a novel workflow designed to time deletions within large chromosomal gains (**Supplementary Fig. 9A-B**) [7]. Overall, we identified only one patient (MM064) with 4p deletions caused by t(4;14)(*NSD2;IGH*), occurring within a large chromosome 4 gain. These deletions consistently caused a copy number jump of 2. Specifically, the t(4;14)(*NSD2;IGH*) deletion resulted in a single allelic copy, while the remaining 4p regions were duplicated (3 vs 1 copies; **Supplemental Data Fig. 1D**). This temporal pattern can be explained by the initial occurrence of t(4;14)(*NSD2*;*IGH*) causing the 4p deletion. Subsequently, the entire deleted allele underwent a large duplication, except for the deleted segments, resulting in a 3:1 CNV jump. Overall, despite the limited number of cases, in particular for *NSD2* and *MAF/MAFB* IGH translocated patients, our data suggest that canonical IGH translocations precede HY and other gains (**Supplemental Data Fig. 1E**).

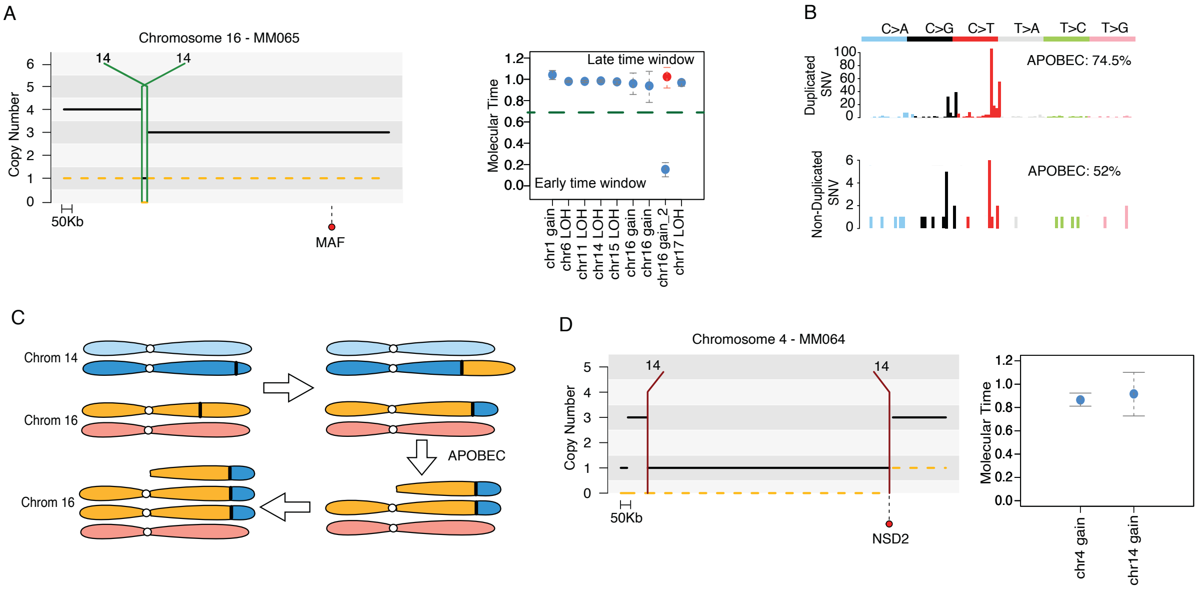
A
Chromosome 16 - MM065
Copy Number
14
14
50Kb
MAF
Molecular Time
Late time window
Early time window
chr1 gain
chr6 LOH
chr11 LOH
chr14 LOH
chr15 LOH
chr16 gain
chr16 gain
chr16 gain_2
chr17 LOH
B
C>A
C>G
C>T
T>A
T>C
T>G
Duplicated SNV
APOBEC: 74.5%
Non-Duplicated SNV
APOBEC: 52%
C
Chrom 14
Chrom 16
APOBEC
Chrom 16
D
Chromosome 4 - MM064
Copy Number
14
14
50Kb
NSD2
Molecular Time
chr4 gain
chr14 gain

**Supplementary Data 4 References**


1 Walker, B. A. *et al.* APOBEC family mutational signatures are associated with poor prognosis translocations in multiple myeloma. *Nat Commun* **6**, 6997 (2015). https://doi.org/10.1038/ncomms7997

2 Rustad, E. H. *et al.* Timing the initiation of multiple myeloma. *Nat Commun* **11**, 1917 (2020). https://doi.org/10.1038/s41467-020-15740-9

3 Maura, F. *et al.* Genomic Classification and Individualized Prognosis in Multiple Myeloma. *J Clin Oncol*, JCO2301277 (2024). https://doi.org/10.1200/JCO.23.01277

4 Samur, M. K. *et al.* Genome-Wide Somatic Alterations in Multiple Myeloma Reveal a Superior Outcome Group. *J Clin Oncol* **38**, 3107-3118 (2020). https://doi.org/10.1200/JCO.20.00461

5 Schavgoulidze, A. *et al.* Prognostic impact of translocation t(14;16) in multiple myeloma according to the presence of additional genetic lesions. *Blood Cancer J* **13**, 160 (2023). https://doi.org/10.1038/s41408-023-00933-4

6 Walker, B. A. *et al.* Identification of novel mutational drivers reveals oncogene dependencies in multiple myeloma. *Blood* **132**, 587-597 (2018). https://doi.org/10.1182/blood-2018-03-840132

7 Cirrincione, A. *et al.* Revealing Novel Mechanisms Underlying Inactivation of Tumor Suppressor Genes on Duplicated Chromosomes in Multiple Myeloma. *Blood* **142**, 874-874 (2023). https://doi.org/10.1182/blood-2023-186802
