## Supplementary Figures for "Temporal Genomic Dynamics Shape Clinical Trajectory in Multiple Myeloma"

**Supplementary Figure 1. The 12 newly diagnosed multiple myeloma (NDMM) genomic groups and their distribution across the two cohorts of NDMM included in this study (Maura et al. JCO 2024).**

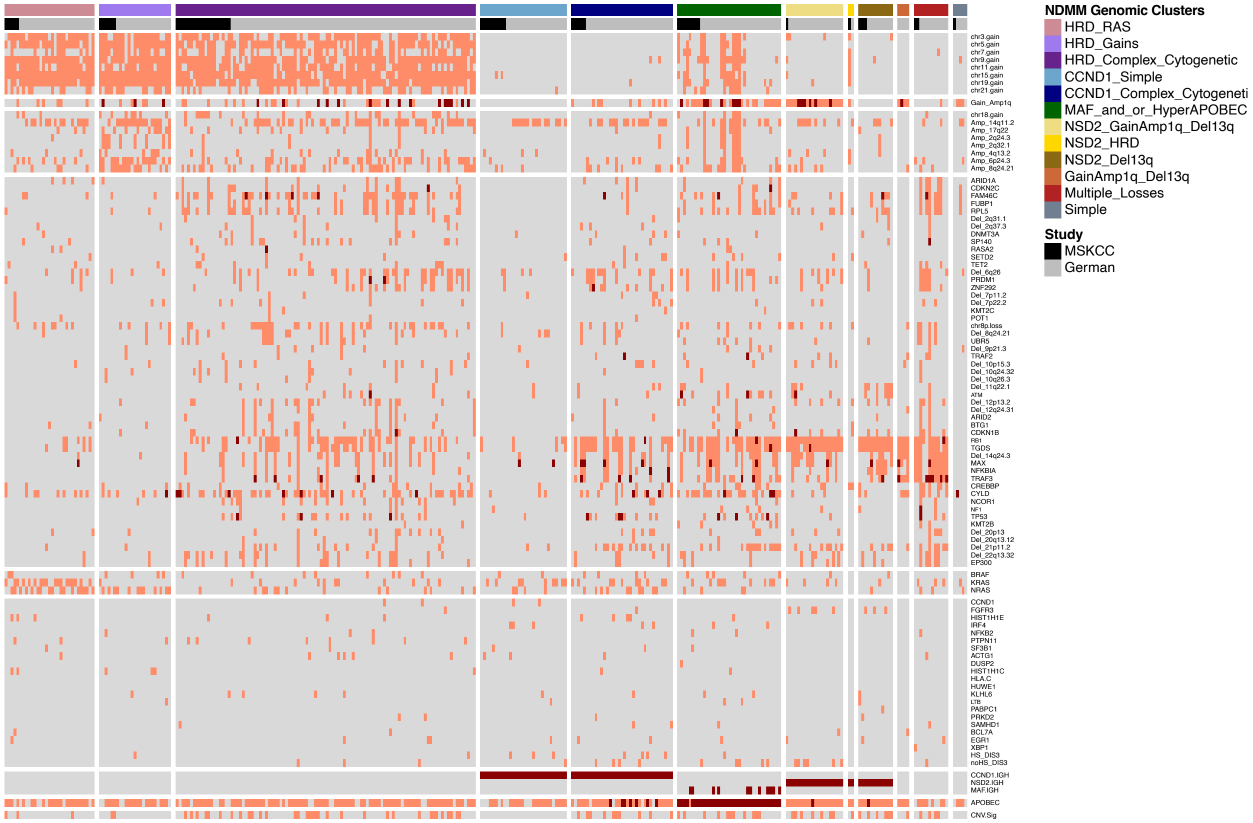

**Supplementary Figure 2. Mutational burden in multiple myeloma. A)** Mutational burden comparison between the two cohorts included in this study. P-value was estimated using Wilcoxon test. **B-C)** Hyper-APOBEC impact on progression free survival (PFS) and overall survival (OS) in newly diagnosed multiple myeloma.

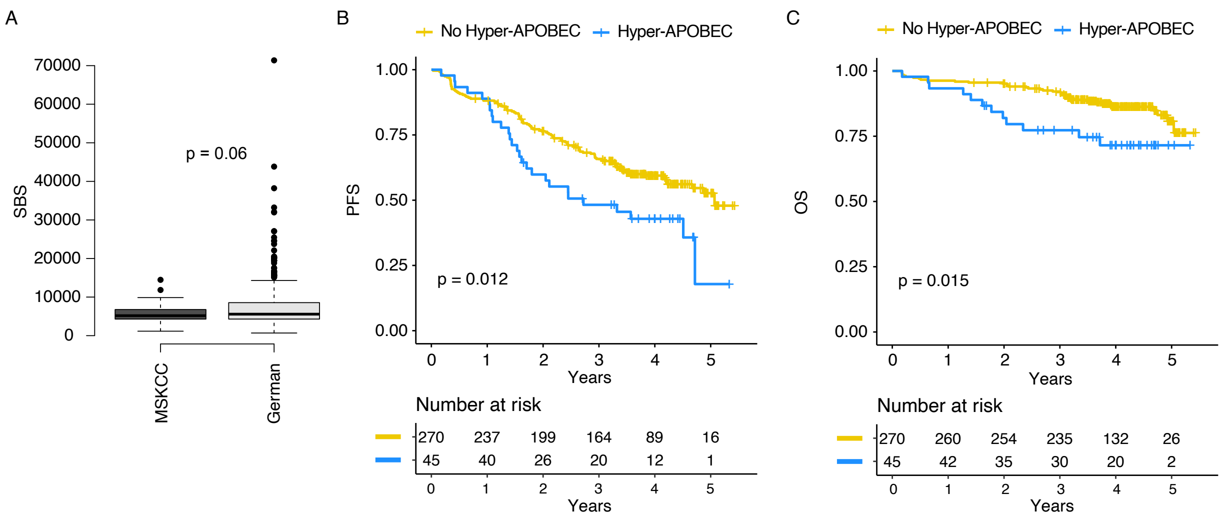

**Supplementary Figure 3. A)** Correlation between SBS1 and SBS5 mutational burden (clock-like signatures) and patients' age at sample collection. Samples that did not fit into the linear model reported in **Fig. 1B** are plotted with more transparent colors (residual >1900). Three samples with more than 8000 SBS1 and SBS5 mutations were not included in the plot for graphical purpose (MM119, MM165, and RRMM48). **B)** Comparison in SBS1 and SBS5 clock mutational rate between multiple myeloma and B-cell lymphoma. **C)** Comparison in SBS1 and SBS5 clock mutational rate between multiple myeloma and normal memory B-cells. Because of the single cell colony expansion behind the normal memory B-cell data (Machado et al. Nature 2022), for the multiple myeloma we considered only the clonal variants. SBS: single base substitution; NDMM: newly diagnosed multiple myeloma; RRMM: relapsed refractory multiple myeloma; Mel+ and Mel-: melphalan exposed or not exposed RRMM patients, respectively.

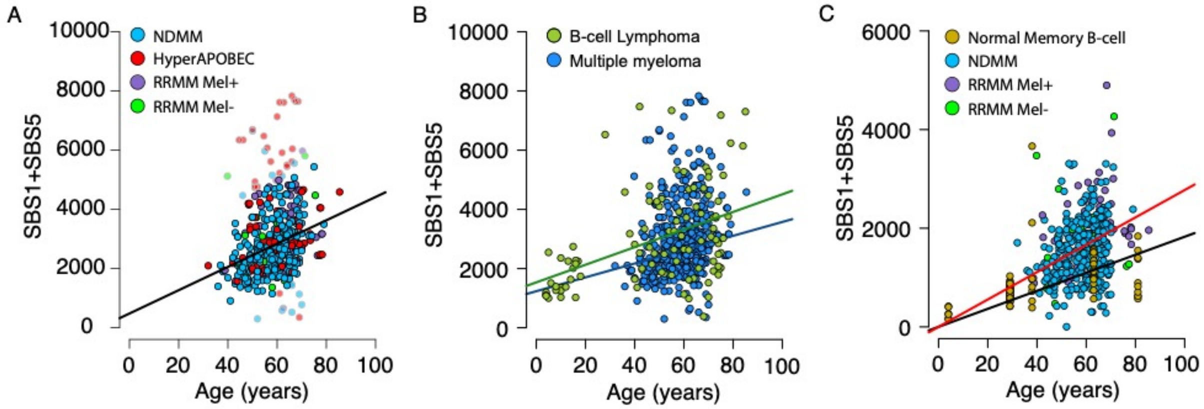

**Supplementary Figure 4. Multiple myeloma pathogenetic model.**

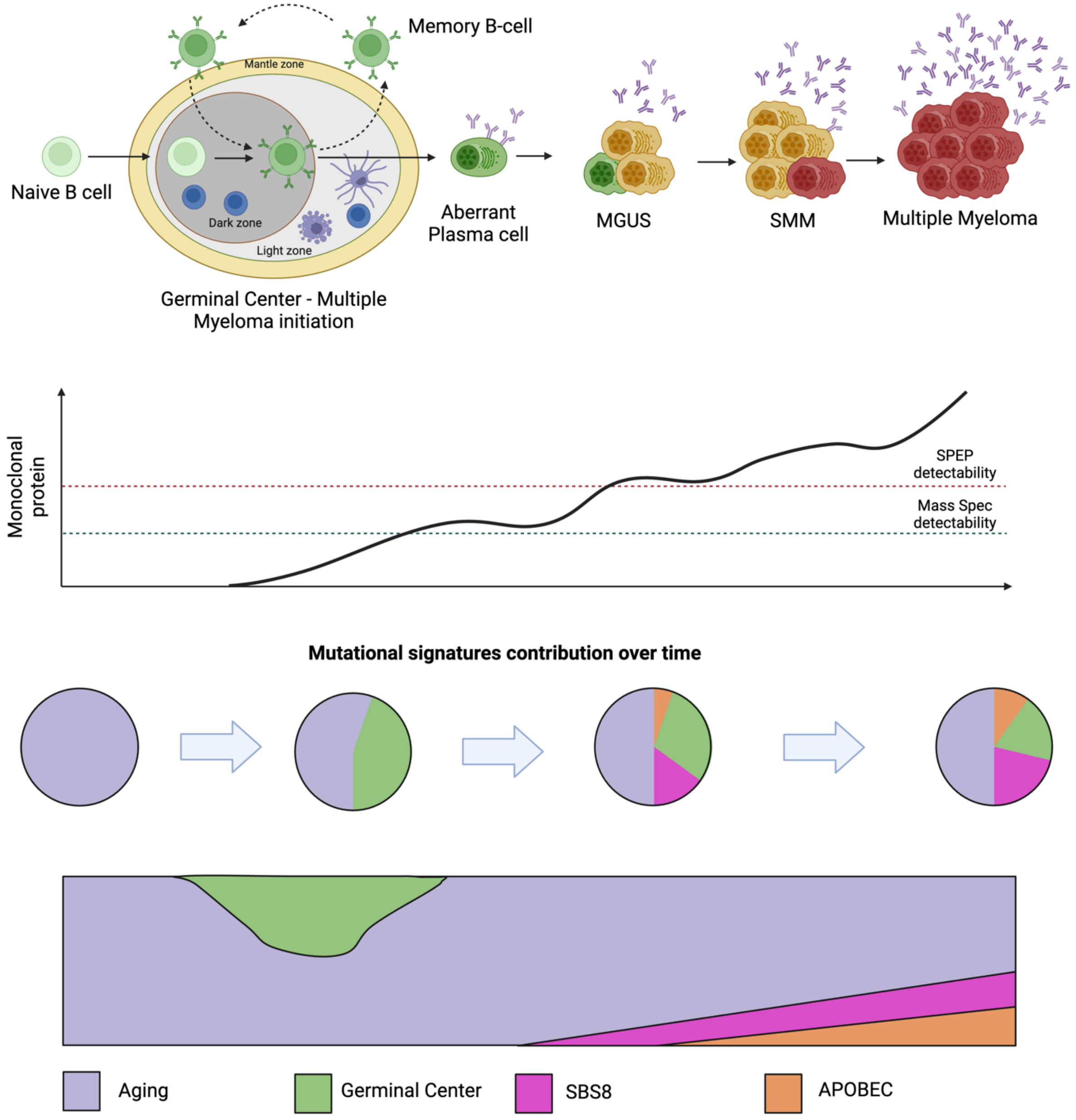

**Supplementary Figure 5.** SBS mutational signature contribution among clonal and subclonal variants among samples with hyper-APOBEC.

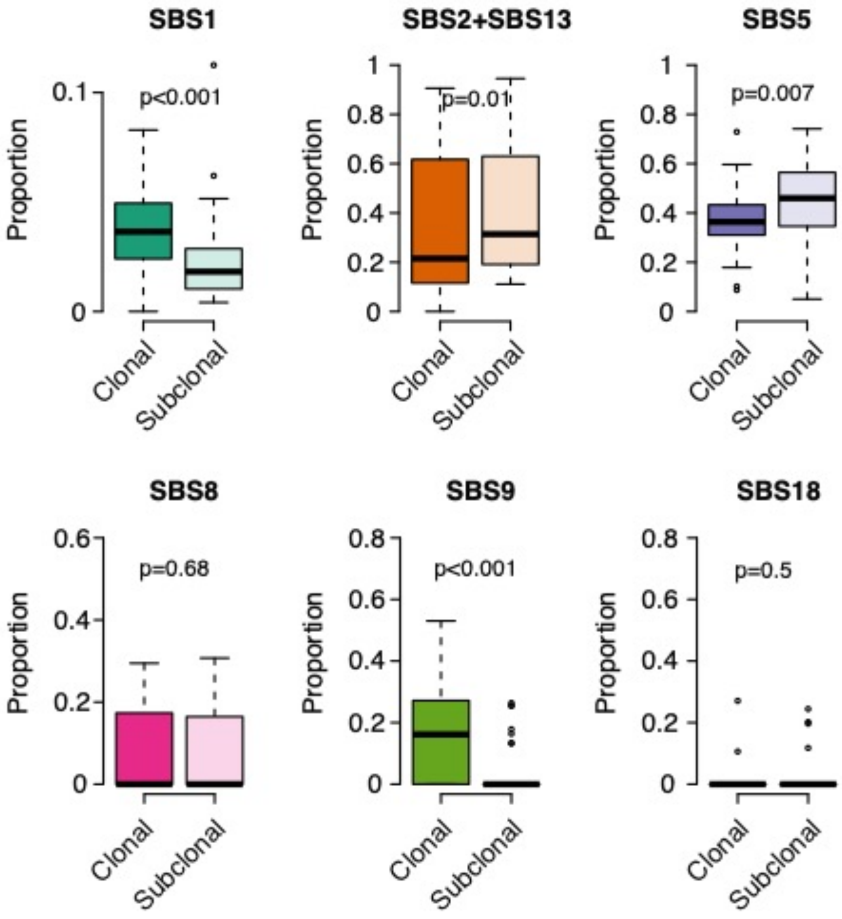

**Supplementary Figure 6. Temporal relationship of distinct genomic subgroups and chromosomes. A)** The molecular time estimates of chromosomal gains that were shared, selected, or lost over time in patients with WGS data available at two different time points was analyzed. **B)** Boxplot of molecular time of non-HY chromosomes (1, 2, 4, 6, 8, 10, 12, 13, 14, 16, 17, 18, 20, and 22) based on the presence (left box) or absence (right box) of canonical IGH translocations. The same pattern is seen among these chromosomes that tend to be acquired later when a canonical IGH translocation is present. **C)** Boxplot of molecular time of odd-numbered, HY chromosomes (3, 5, 7, 9, 11, 15, 19, and 21) (y axis) based on presence (left box) or absence (right box) of canonical IGH translocations (x axis). Overall, the co-occurrence of a canonical IGH translocation was associated with a later molecular time of HY chromosomal gains. **D)** Boxplot of molecular time of all CN-LOH among HY patients, divided into odd-numbered HY chromosomes (left box), and non-HY chromosomes (right box). All p-values were estimated using Wilcoxon test.

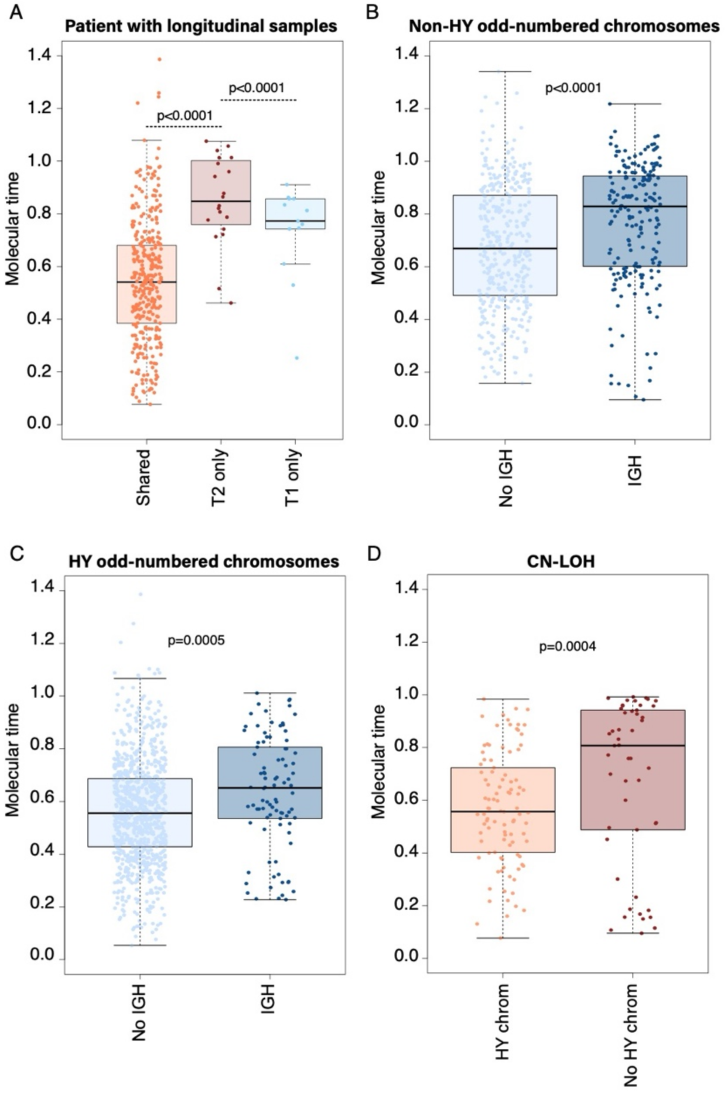

**Supplementary Figure 7. Temporal patterns of gains across multiple myeloma 12 genomic groups (Maura et al. JCO 2024). A)** Molecular time (y axis) of all chromosomal duplications across the 12 genomic groups considering all patients. P-adjusted values for each comparison are reported in **Supplementary Table 8**. **B)** Molecular time (y axis) of all odd-numbered HY chromosomal duplications across the 12 genomic groups. Only patients with HY are reported. P-adjusted values for each comparison are reported in **Supplementary Table 8**. **C)** Molecular time (y axis) of all 1q gains across the 12 genomic groups. P-adjusted values for each comparison are reported in **Supplementary Table 8**.

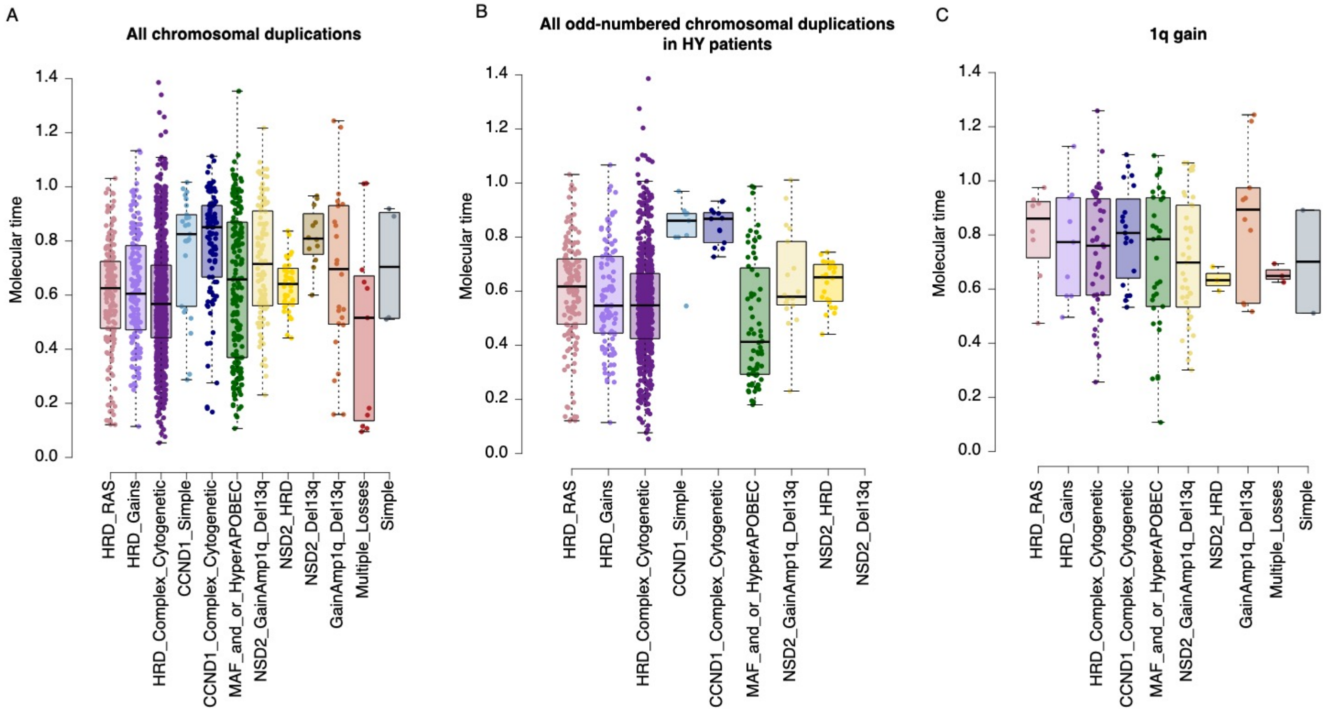

**Supplementary Figure 8. Timing focal deletions and gains such as MYC in multiple myeloma.** Cartoon summarizing the pre- (**A**) and post-gain (**B**) deletion workflow, and the pre- (**C**) and post-duplication (**D**). In **E** and **F** we provided two possible temporal scenarios where MYC translocation can be acquired after (**E**) or before large chromosomal gains (**F**).

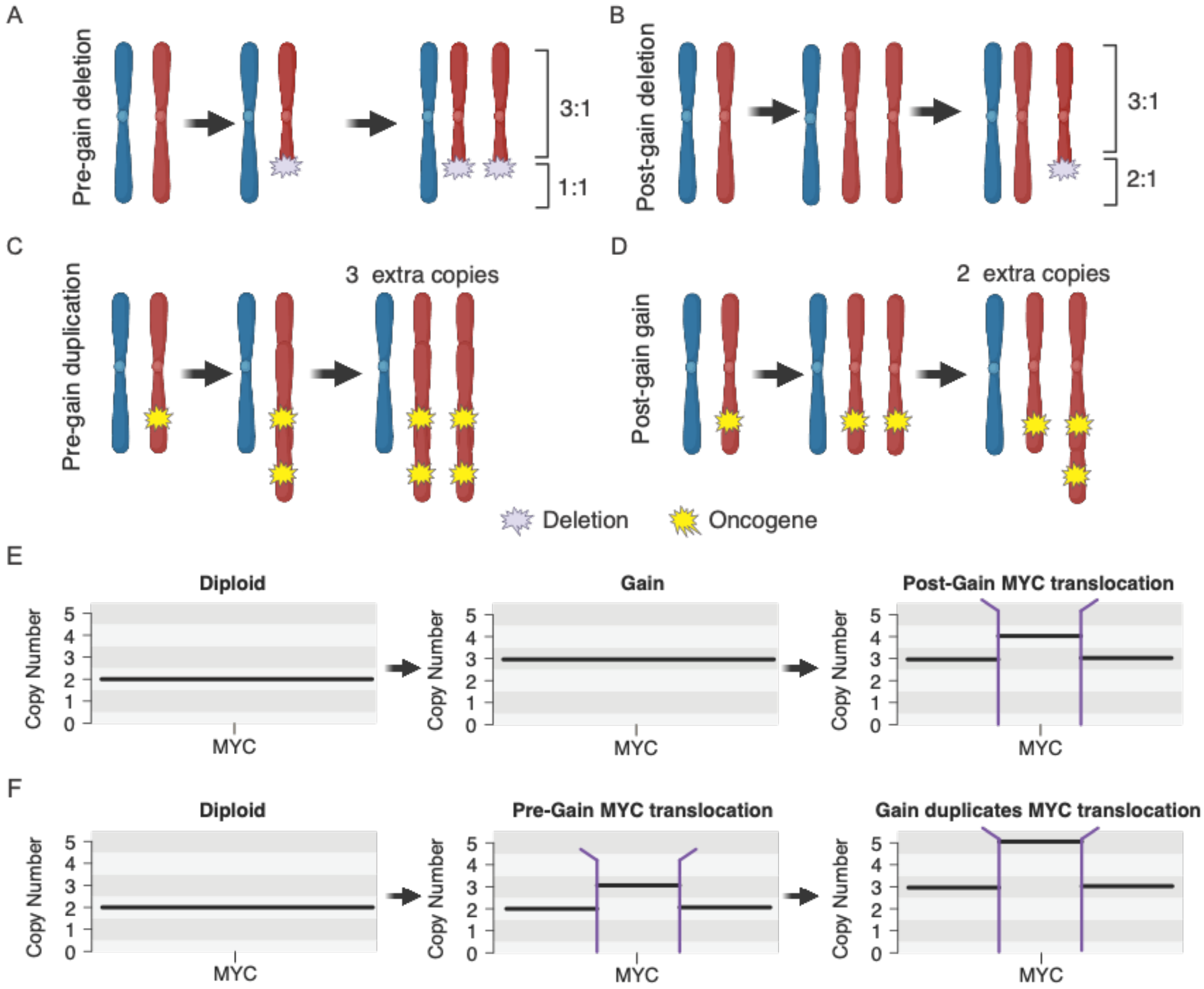
